# EpiZoo: a DNA sequence-aware foundation model for cross-species single-cell epigenomics

**DOI:** 10.64898/2026.09.24.754017

**Authors:** Keyi Li, Xiaoyang Chen, Qun Jiang, Zian Wang, Hairong Lv, Rui Jiang

## Abstract

Large-scale single-cell epigenomic atlases characterize chromatin regulatory landscapes across diverse biological contexts, spanning cell types, tissues, individuals and species. Foundation models provide an opportunity to capture the full spectrum of cellular diversity in these atlases, yet current models remain largely confined to individual species by genomic coordinate dependence and overlook regulatory information encoded in DNA sequences. Here we introduce EpiZoo, a DNA sequence-aware foundation model for cross-species single-cell epigenomics. EpiZoo converts million-dimensional single-cell epigenomic profiles from diverse species into compact cell sentences that integrate DNA-encoded regulatory information, sequence-independent epigenomic context and accessibility-based importance. Built around a mixture-of-experts transformer and containing 2.6 billion parameters in total, EpiZoo is pretrained on our manually curated multi-species Omni-scATAC corpus of approximately 20.9 million cells to learn regulatory programs across species. On external datasets excluded from pretraining, EpiZoo achieves state-of-the-art performance in fundamental single-cell analysis tasks, including feature extraction, cell type annotation and data imputation. Its sequence-aware architecture enables extension to evolutionarily diverse species, and supports comparative analysis of regulatory conservation and divergence during primate evolution. Benefiting from this modeling design, EpiZoo enables context-aware prioritization of somatic mutations in cancer and prediction of cell-type-specific chromatin accessibility from DNA sequences across genomic regions and species.

## Introduction

Epigenomic regulation provides a critical molecular interface between genomic sequence and cellular phenotype by controlling *cis*-regulatory element activity and transcriptional programs^1,2^. Single-cell epigenomic profiling technologies, particularly single-cell assay for transposase-accessible chromatin using sequencing (scATAC-seq)^3^, enable these regulatory programs to be characterized at single-cell resolution^4^. High-throughput scATAC-seq studies have accelerated the generation of large-scale single-cell epigenomic atlases across diverse biological contexts, providing valuable resources for investigating how candidate *cis*-regulatory elements (cCREs) contribute to cell identity, lineage specification and disease-associated regulatory programs^5-7^. With increasing coverage across species, these resources enable comparisons of regulatory landscapes across evolutionary distances^8^, identification of conserved and species-specific cCRE co-accessibility patterns^9,10^, and investigation of how DNA sequence variation influences epigenomic states and cellular phenotypes^11^. However, the extreme sparsity and high dimensionality of scATAC-seq data pose substantial challenges for direct downstream analysis^12,13^.

To address these analytical challenges, numerous methods have been developed for scATAC-seq analysis^13-18^. However, because most of them are designed for specific tasks and optimized within individual datasets, the patterns they capture can remain associated with tissue contexts or cell-type compositions, rather than reflecting generalizable regulatory information underlying single-cell epigenomic data^18,19^. As a result, these methods do not fully exploit the epigenomic landscapes profiled in large-scale scATAC-seq atlases and often exhibit unstable performance and limited generalizability when applied to new datasets. Moreover, extending them to new tasks typically requires substantial methodological modifications or complete model retraining. These limitations hinder the reuse of the extracted information across datasets and tasks, underscoring the need for a scalable and unified framework for modeling single-cell regulatory landscapes.

Foundation models have recently emerged for single-cell epigenomics, aiming to capture cellular heterogeneity and regulatory programs from scATAC-seq atlases^20-22^. As a representative model, EpiAgent has demonstrated strong performance across fundamental downstream tasks and showed the potential of large-scale pretraining for expanding the analytical utility of single-cell epigenomic data^20^. However, existing foundation models remain constrained by two major limitations. First, their restriction to single species limits the scale and heterogeneity of pretraining data, prevents cross-species transfer and joint analysis, and hampers the discovery of conserved and species-specific regulatory programs^8,23^. Second, these models do not explicitly incorporate DNA sequence information, preventing them from capturing the sequence grammar that links nucleotide variation to chromatin accessibility and assessing single-nucleotide effects in cell-type or disease contexts^11,24-26^. Together, these limitations demand a foundation model that enables cross-species regulatory analyses beyond species-specific genomic coordinates, and connects DNA sequence variation with chromatin accessibility and phenotypic effects through single-cell epigenomic data.

To fill this gap, we introduce EpiZoo, a DNA sequence-aware foundation model for cross-species single-cell epigenomics. EpiZoo tokenizes each cell by converting highly sparse chromatin accessibility profiles spanning over one million cCREs into compact cell sentences containing at most 8,192 tokens. This tokenization strategy reduces input dimensionality while retaining informative regulatory signals. Instead of relying solely on species-specific genomic coordinates, EpiZoo uses a sequence-to-embedding anchoring module (SEAM) to encode the underlying DNA sequences of cCREs into sequence-derived regulatory priors. These sequence embeddings are combined with learnable identity and rank embeddings for the corresponding cCRE tokens to form sequence-aware token embeddings, which are then processed by a mixture-of-experts (MoE)^27^ transformer to capture long-range co-accessibility patterns across species.

Extensive benchmarking demonstrates that EpiZoo consistently outperforms existing methods across fundamental downstream tasks for single-cell analysis, including feature extraction, cell type annotation and data imputation. Our evaluations also identified a stable and efficient fine-tuning strategy, in which low-rank adaptation (LoRA) preserves pretrained modeling priors and avoids overfitting to dataset-specific contexts observed with full-parameter fine-tuning (FFT). More importantly, the DNA sequence-aware design of EpiZoo extends single-cell epigenomic modeling beyond species-specific genomic coordinates. By using SEAM to encode transferable regulatory priors, EpiZoo can be post-trained on small-scale datasets from previously unseen species to establish species-specific and cross-species foundation models. Building on this cross-species capacity, EpiZoo-Evo, an EpiZoo extension for joint primate modeling, reveals continuous gradients of regulatory conservation and divergence in primate neurodevelopment, and identifies sequence-divergent cCREs with putatively shared functions. The DNA sequence-aware architecture further supports nucleotide-level inference. EpiZoo-Cancer, an EpiZoo extension another extension of EpiZoo for cancer contexts, prioritizes noncoding somatic mutations and refines their predicted effects in diseases. By concatenating cell-type embeddings and sequence embeddings, EpiZoo also predicts cell-type-specific chromatin accessibility from DNA sequence, reconstructs continuous chromatin accessibility profiles across unseen regions and identifies candidate motifs underlying enhancer activity.

## Results

### EpiZoo is a DNA sequence-aware foundation model for cross-species single-cell epigenomics

EpiZoo is a large-scale foundation model designed to capture regulatory programs underlying single-cell chromatin accessibility profiles across species (Fig. 1). The model comprises a DNA sequence-aware embedding module, an MoE transformer and species-specific signal decoders, and is pretrained on our manually curated Omni-scATAC corpus using two complementary species-specific tasks. This design supports extensions to cross-species analysis and DNA sequence-based inference under different contexts.

**Fig. 1:**
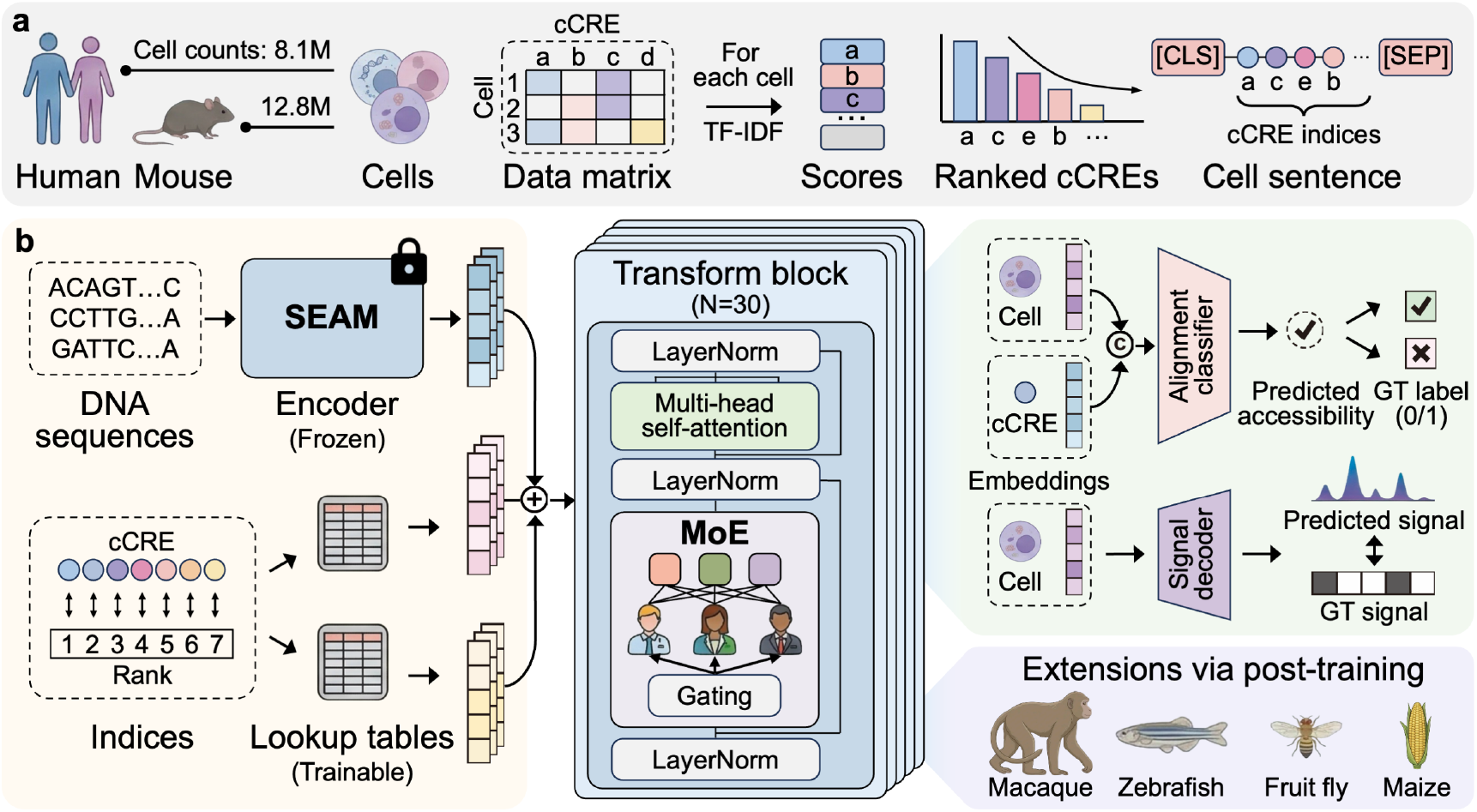
EpiZoo is a DNA sequence-aware foundation model for cross-species single-cell epigenomics. **a**, Omni-scATAC construction and cell sentence tokenization. EpiZoo is pretrained on our manually curated human-mouse scATAC-seq corpus, namely Omni-scATAC, comprising approximately 20.9 million cells across more than 30 tissues and cell lines. Each sscell is converted into a cell sentence composed of a [CLS] token, TF-IDF-ranked accessible cCRE tokens and a [SEP] token. **b**, Model architecture, pretraining strategy and cross-species extension. EpiZoo consists of a DNA sequence-aware embedding module, an MoE transformer and species-specific signal decoders. The model is pretrained using two complementary tasks, including cell-cCRE alignment and signal reconstruction, and can be extended to new species via post-training.

We manually curated Omni-scATAC, a large-scale scATAC-seq corpus of 46 public human and mouse datasets, covering approximately 20.9 million cells across more than 30 tissues and cell lines (Fig. 1a and Supplementary Table 1). To support scalable pretraining across these heterogeneous datasets, we processed each profile into a standardized cell-by-cCRE matrix in the corresponding species-specific cCRE feature space. EpiZoo then converts each cell into a cell sentence composed of a [CLS] token, accessible cCRE tokens, and a [SEP] token, with the total length capped at 8,192 tokens. Specifically, accessible cCREs are ranked in descending order by term frequency-inverse document frequency (TF-IDF) scores. This tokenization strategy converts the extremely sparse and high-dimensional scATAC-seq data into compact cell sentences while preserving both the relative importance of accessible cCREs and their regulatory information in each cell.

EpiZoo contains approximately 2.6 billion parameters and consists of three core modules: a DNA sequence-aware embedding module, an MoE transformer and species-specific signal decoders (Fig. 1b and Supplementary Table 2). The embedding module converts input cell sentences into token embeddings by combining three complementary components for each cCRE token: a sequence embedding generated by SEAM, a learnable identity embedding and a learnable rank embedding. SEAM encodes the underlying DNA sequence of each cCRE to provide regulatory priors and reduce dependence on species-specific genomic coordinates. In parallel, the learnable identity embedding captures sequence-independent epigenomic context, whereas the rank embedding encodes the importance of each accessible cCRE according to its TF-IDF ranking. The resulting token embeddings are then processed by the MoE transformer, in which sparse expert routing captures long-range co-accessibility patterns while allowing different experts to specialize in regulatory heterogeneity associated with species, tissues and cell types. The output [CLS] token embedding is used as the cell embedding, summarizing the heterogeneity of each cell at the epigenomic level. Finally, species-specific signal decoders map each cell embedding to the corresponding cCRE feature space to reconstruct the global chromatin accessibility.

EpiZoo is pretrained using a multi-task framework comprising cell-cCRE alignment and signal reconstruction within species-specific cCRE vocabularies. In the cell-cCRE alignment task, the model concatenates each cell embedding with identity embeddings of a sampled cCRE set to predict whether each cCRE in the set is accessible in the corresponding cell. In the signal reconstruction task, the species-specific signal decoders reconstruct the global chromatin accessibility profile from the cell embedding alone. During early pretraining, we periodically calibrated SEAM with identity embeddings to align it with the model embedding space and to encourage the extraction of DNA-related regulatory priors from scATAC-seq data. SEAM was then frozen for all subsequent pretraining stages, downstream fine-tuning and applications. Through pretraining on the Omni-scATAC corpus, EpiZoo learns generalizable patterns that support efficient fine-tuning for downstream analyses and post-training for phylogenetically diverse species with limited data. Details of data curation, model architecture and pretraining framework are provided in Methods.

### EpiZoo facilitates fundamental downstream tasks for single-cell analysis

The extreme sparsity and high dimensionality of scATAC-seq data present substantial challenges for direct downstream analyses, making reliable low-dimensional cell embeddings essential for resolving cellular heterogeneity and supporting subsequent analytical tasks^12^. Here, we evaluated EpiZoo across three fundamental downstream tasks centered on these embeddings: feature extraction, cell type annotation and data imputation. The benchmarks were conducted on four representative datasets excluded from pretraining, including Kanemaru2023 (human heart)^28^, Li2023b (human brain)^29^, Cusanovich2018 (multiple mouse tissues)^30^, and Fang2021 (mouse brain)^13^.

### EpiZoo generates embeddings that capture cellular heterogeneity

Effective feature extraction should reduce the dimensionality of sparse chromatin accessibility profiles while retaining biological variation that distinguishes cell populations, thereby providing a basis for subsequent scATAC-seq analyses. We used unsupervised Louvain clustering to benchmark the quality of cell embeddings generated by EpiZoo and six baseline methods, including EpiAgent^20^, CASTLE^31^, cisTopic^17^, scBasset^32^, PeakVI^33^, and SCALE^34^.

After Fine-tuning, EpiZoo consistently outperforms baseline methods in cell clustering, as measured by both normalized mutual information (NMI) and adjusted rand index (ARI) scores (Fig. 2a and Supplementary Figure 1). This improvement is observed across benchmark datasets spanning diverse human and mouse tissues, indicating that EpiZoo cell embeddings preserve regulatory patterns that is more informative for resolving cellular heterogeneity under different biological contexts. Notably, zero-shot EpiZoo also achieves competitive performance on these held-out datasets excluded from pretraining, suggesting that the model architecture and the pretraining stage allow EpiZoo to learn transferable and generalizable patterns before dataset-specific fine-tuning. Embedding ablation analysis further shows that the full EpiZoo model outperforms DNA sequence-only and cCRE identity-only variants, highlighting the complementary and indispensable roles of DNA sequence information and sequence-independent epigenomic context (Supplementary Note 1 and Supplementary Figures 2 and 3). Consistently, UMAP visualizations show that EpiZoo cell embeddings form more compact clusters within each cell type and better separate closely related cell populations (Fig. 2b). For example, in the Kanemaru2023 dataset^28^, adipocytes and fibroblasts are related stromal populations with overlapping mesenchymal regulatory programs. EpiZoo distinguishes these two cell types that remained partially conflated by baseline methods, indicating improved resolution of fine-grained cellular heterogeneity.

**Fig. 2:**
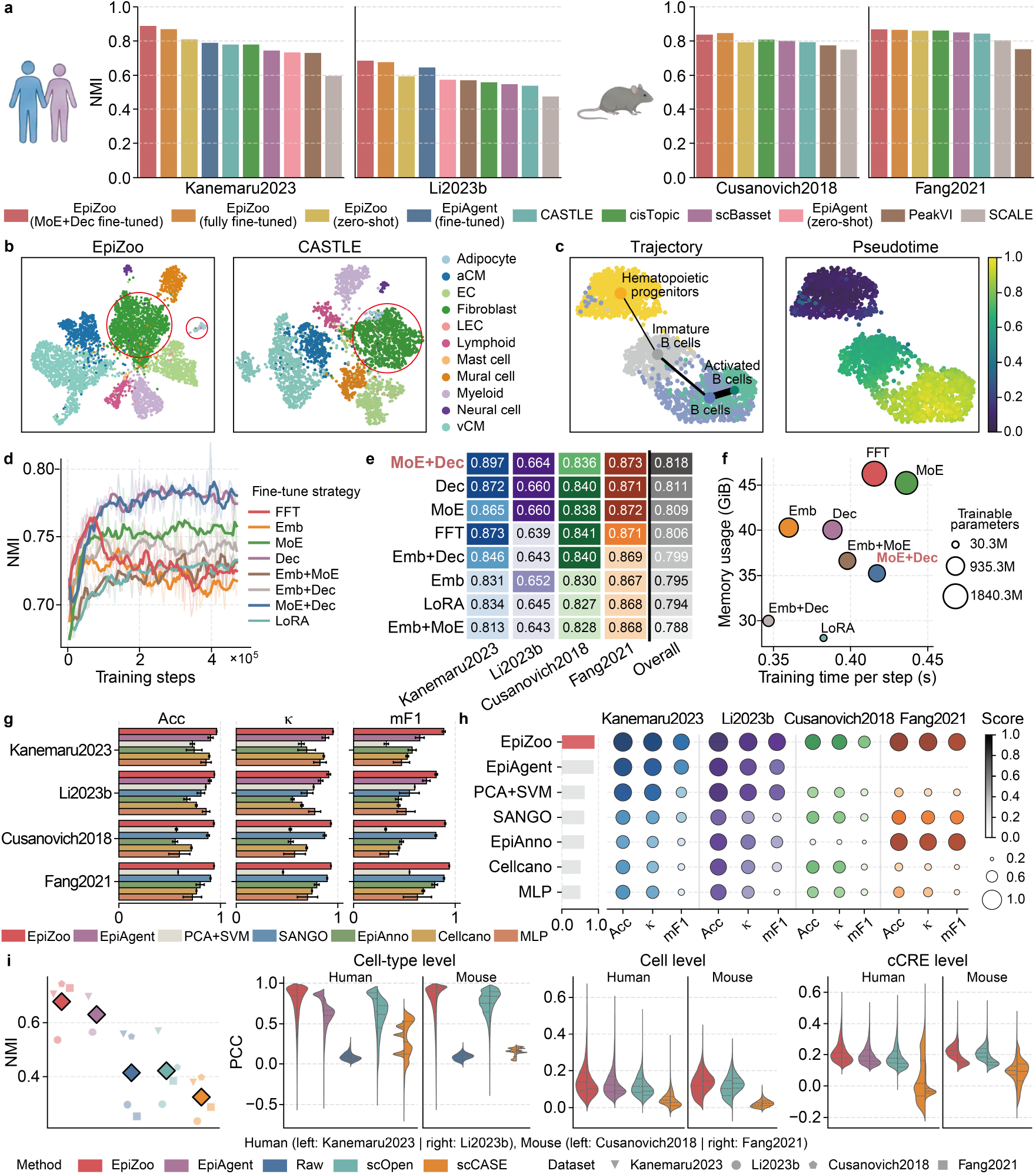
EpiZoo facilitates fundamental downstream tasks for single-cell analysis. **a**, Benchmarking performance on feature extraction across four held-out datasets, assessed by NMI following Louvain clustering. **b**, UMAP visualization of cell embeddings from EpiZoo (left) and CASTLE (right) on the Kanemaru2023 dataset. Circled regions indicate fibroblast and adipocyte populations. **c**, PAGA-inferred B-cell developmental trajectory (left) and DPT values (right) based on EpiZoo cell embeddings in the Cusanovich2018 dataset. Edge width in the trajectory plot indicates the PAGA-inferred connectivity strength between cell populations. **d**, Step-wise performance of different fine-tuning strategies, measured by NMI following Louvain clustering. Light-colored lines indicate step-wise NMI scores, and dark-colored lines indicate NMI trends smoothed with a five-step moving window. **e**, Benchmarking performance of different fine-tuning strategies, measured by NMI following Louvain clustering. Overall denotes the average performance of each strategy across the four benchmark datasets. **f**, Computational cost of different fine-tuning strategies, measured by GPU memory usage and training time. For **d-f**, FFT indicates full-parameter fine-tuning of all three modules, and LoRA indicates low-rank adaptation applied to all modules. Emb, MoE and Dec denote LoRA applied to the DNA sequence-aware embedding module, MoE transformer and signal decoders, respectively; unlisted modules were updated by FFT. **g**, Benchmarking performance on cell type annotation under intra-dataset five-fold cross-validation, evaluated by accuracy (Acc),κ and mF1. Bar heights indicate mean values, and error bars indicate 95% confidence intervals. **h**, Benchmarking performance on cell type annotation under inter-dataset validation, evaluated by Acc, κ and mF1. **i**, Benchmarking performance on data imputation using datasets with 50% simulated random dropout. From left to right, panels show NMI scores after Louvain clustering (left) and Pearson correlation coefficients (PCCs) between imputed and reference matrices at the cell-type (middle left), cell (middle right) and cCRE (right) levels. The inner horizontal lines in the violin plots denote the first quartile, median, and third quartile.

Beyond resolving discrete cell populations, EpiZoo embeddings also reveal continuous lineage progression. We first generated EpiZoo embeddings for B-lineage cells from the Cusanovich2018 dataset^30^, and then PAGA-based trajectory inference^35^ and diffusion pseudotime (DPT) analysis to these embeddings. As shown in Fig. 2c, the resulting trajectories clearly delineate a hierarchical differentiation path from hematopoietic progenitors to immature B cells, mature B cells and activated B cells, and DPT values resolve cell-level progression along this continuum. This ordering is consistent with the expected differentiation process of B-lineage cells. Together, these results indicate that EpiZoo embeddings capture both discrete cell types and continuous developmental heterogeneity.

When fine-tuning the large-scale EpiZoo model on smaller downstream datasets, unconstrained FFT can overfit dataset-specific chromatin accessibility patterns and degrade pretrained regulatory information, as reflected by a rapid decline in cell embedding quality after the initial training steps (Fig. 2d and Supplementary Figure 4). To identify a more stable and efficient fine-tuning strategy, we evaluated eight configurations in which the embedding module, MoE transformer and signal decoders were each adapted either by FFT or by LoRA^36^. The best-performing configuration applies FFT to the embedding module and LoRA to the MoE transformer and signal decoders. Compared with the standard FFT, this hybrid strategy shows more stable performance gains during training (Fig. 2d), achieves comparable or superior cell embedding quality across configurations as assessed by Louvain clustering (Fig. 2e), and reduces both GPU memory usage and training time (Fig. 2f).

### EpiZoo enables precise cell type annotation

Manual cell type annotation in scATAC-seq data remains complex, labor-intensive, and difficult to standardize across studies. By learning embeddings that resolve cellular heterogeneity, EpiZoo can be fine-tuned on labeled reference datasets and then applied to previously unseen scATAC-seq profiles for accurate and scalable cell type annotation.

We first evaluated EpiZoo using intra-dataset five-fold cross-validation (Fig. 2g and Supplementary Figures 5-8). EpiZoo consistently outperforms all evaluated baseline methods^37-41^ across the four datasets, while maintaining high annotation accuracy in datasets comprising multiple tissues or closely related cell populations. On the Cusanovich2018 dataset^30^, a multi-tissue mouse atlas containing 39 cell types with uneven numbers of cells, EpiZoo achieves an annotation accuracy of 0.935 and a macro F1 score (mF1) of 0.901. In the human brain Li2023b and mouse brain Fang2021 datasets, baseline methods frequently confuse closely related neuronal populations, such as excitatory and inhibitory subtypes or neurons from distinct cortical layers. In contrast, EpiZoo more accurately resolves these fine-grained populations and assigns neuronal cells to their correct subtype labels.

We further evaluated EpiZoo under a more challenging and practical scenario, namely inter-dataset annotation (Fig. 2h and Supplementary Figures 9-12). For each held-out benchmark dataset, the model was fine-tuned on a tissue-matched reference dataset from the Omni-scATAC corpus, as detailed in Methods. Across the four datasets, EpiZoo consistently outperforms the baseline methods in accuracy, Cohen’s kappa coefficient (κ) and mF1, with average relative improvements of 9.96%, 12.40% and 23.17%, respectively, calculated against the strongest baseline for each dataset and metric. The larger gains in κ and mF1 further suggest that EpiZoo more effectively captures cell-type-specific chromatin accessibility patterns, thereby better distinguishing less abundant or closely related cell populations. Compared with intra-dataset validation, most baseline methods show performance degradation under inter-dataset validation, reflecting their sensitivity to dataset-specific technical variation. In contrast, EpiZoo limits the decrease in accuracy to 0.053 in the Kanemaru2023 dataset and 0.069 in the Fang2021 dataset, whereas the corresponding average declines among baseline methods are 0.111 and 0.216, respectively. These results indicate that EpiZoo’s architecture and pretraining strategy enable robust cell type annotation of independently generated scATAC-seq datasets.

### EpiZoo achieves high-fidelity data imputation

Dropout events in scATAC-seq data introduce substantial noise and exacerbate the intrinsic sparsity of chromatin accessibility profiles, thereby compromising downstream analysis accuracy. After task-specific fine-tuning, EpiZoo projects cell embeddings back to the original high-dimensional cCRE feature space, enabling recovery of dropout events caused by technical sequencing limitations. To evaluate this reconstruction capacity, we introduced simulated dropouts by randomly masking 50% of the non-zero signals in the original profiles and benchmarked EpiZoo against established imputation methods.

To assess how effectively different methods enhance downstream analysis, we first performed cell clustering as a representative evaluation. As shown in Fig. 2i and Supplementary Figure 13, EpiZoo achieves the highest NMI scores across all datasets, indicating more faithful recovery of chromatin accessibility signals that distinguish cell populations after simulated dropout. Consistently, low-dimensional representations derived from EpiZoo-imputed matrices exhibit clearer separation between cell populations compared with other methods (Supplementary Figure 14). Moreover, genomic regions enrichment of annotations tool (GREAT)^42^ analysis in the Kanemaru2023 dataset shows that the imputed signals generated by EpiZoo yield stronger enrichment of cell-type-relevant biological processes than the raw data (Supplementary Note 2 and Supplementary Figure 15). We further evaluated imputation fidelity at the cell-type, cell, and cCRE levels, with Pearson correlation coefficients (PCCs) computed against reference profiles aggregated by cell type, for individual cell and for individual cCRE, respectively. EpiZoo consistently achieves the highest PCCs across all three levels. At the cell-type level, each imputed cell was compared with the aggregated accessibility profile from cells of the same cell type, EpiZoo reaches a median correlation of 0.874, outperforming established methods including EpiAgent^20^, scOpen^43^ and scCASE^44^. These results demonstrate that EpiZoo captures informative regulatory patterns from dropout-affected scATAC-seq data, enabling high-fidelity imputation and enhancement of chromatin accessibility signals.

### EpiZoo empowers foundational cell representations across diverse species

Developing foundation models for non-model organisms is severely bottlenecked by the scarcity of large-scale single-cell epigenomic data. Furthermore, transferring prior knowledge across species in epigenomics is particularly challenging because cCREs are defined by species-specific genome coordinates and often undergo rapid sequence turnover during evolution. Unlike genes or proteins, which can be linked across species through homologous annotations, cCREs lack a universal coordinate system, limiting the ability of existing foundation models to generalize learned regulatory logic to new biological systems. To overcome this barrier, we developed a post-training framework that adapts EpiZoo to construct specific foundation models for unseen species. For each target species, SEAM first encodes the DNA sequence of cCREs into sequence embeddings, providing a functional prior independent of genomic coordinates. The learnable cCRE identity embeddings are then initialized according to evolutionary distance. For phylogenetically proximal species, cCREs that can be mapped to reference cCREs by coordinate lifting inherit the corresponding pretrained cCRE identity embeddings and decoder parameters, whereas unmapped cCREs are randomly initialized. For evolutionarily distant species lacking reliable correspondence, the entire cCRE vocabulary is initialized de novo. We validated the efficacy of this framework across four phylogenetically diverse species: macaque (*Macaca fascicularis*)^45^, zebrafish (*Danio rerio*)^8^, fruit fly (*Drosophila melanogaster*)^8^, and maize (*Zea mays*)^46^ (Fig. 3a and Supplementary Table 3). Comprehensive descriptions of the adaptation strategy, including model initialization and post-training framework are provided in Methods.

**Fig. 3:**
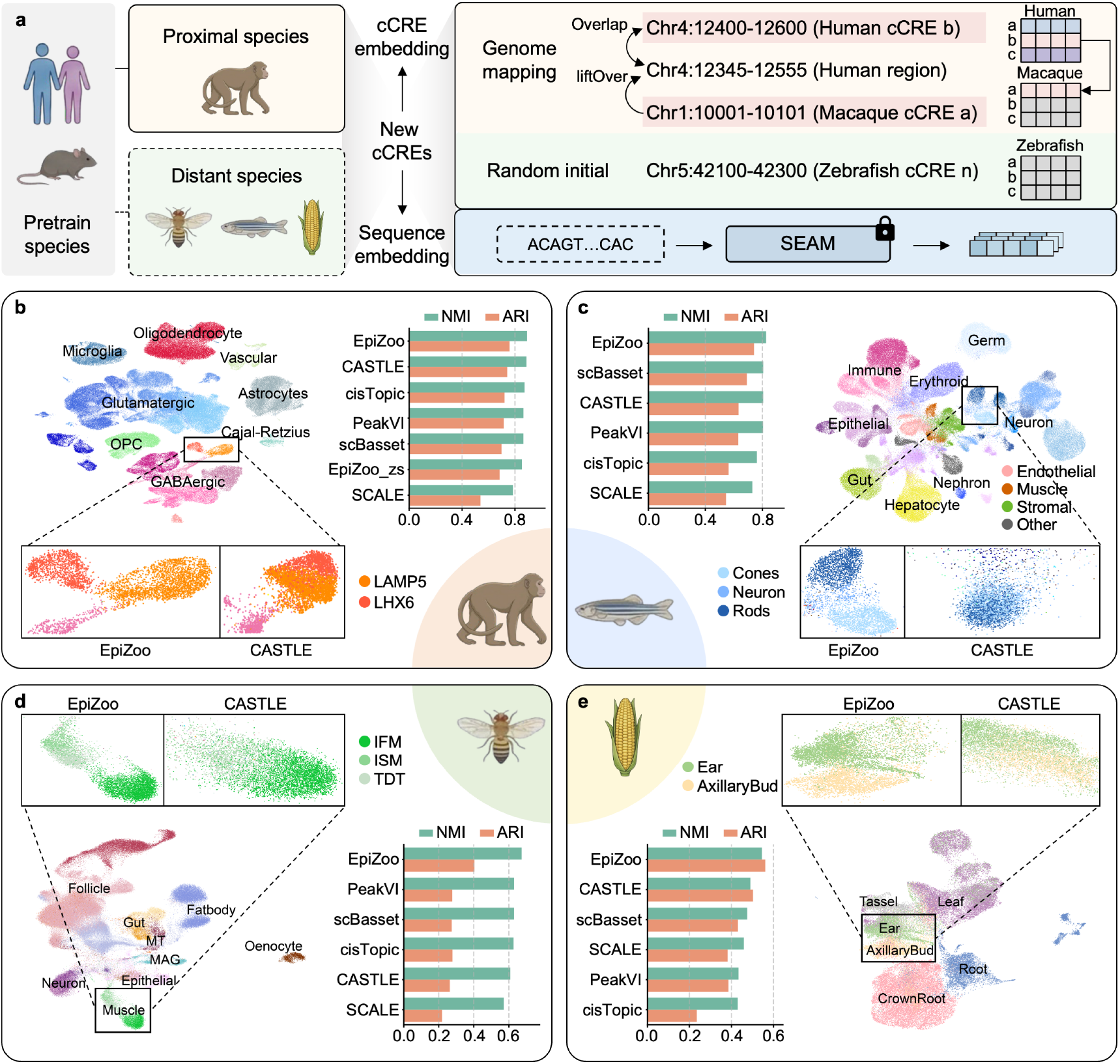
EpiZoo empowers foundational cell representations across diverse species. **a**, Cross-species adaptation strategy according to evolutionary distance. For phylogenetically proximal species, target cCREs are mapped to reference genome using liftOver. Mapped cCREs inherit pretrained embedding and decoder weights, whereas unmapped cCREs are randomly initialized. For evolutionarily distant species, the entire cCRE vocabulary is randomly initialized. For all target species, cCRE DNA sequences are encoded by SEAM. **b**-**e**, Evaluation of species-specific EpiZoo models on macaque (**b**), zebrafish (**c**), fruit fly (**d**) and maize (**e**). UMAP visualizations of EpiZoo cell embeddings labeled by cell types, with zoomed-in regions highlighting closely related populations that are difficult to distinguish. Quality of low-dimensional cell embeddings is compared with baseline methods using NMI and ARI following Louvain clustering. EpiZoo_zs denotes the zero-shot inference using only orthologous cCREs with inherited pretrained weights.

We benchmarked EpiZoo against five baseline methods using cell clustering as a downstream evaluation of embedding quality (Fig. 3b-e). Given the close evolutionary relationship between human and macaque, a substantial proportion of cCREs can be mapped between the two species. We therefore evaluated a zero-shot setting, denoted EpiZoo_zs, where macaque cells were represented using only mapped cCREs, with the corresponding human pretrained parameters directly transferred without macaque-specific post-training. Under this setting, EpiZoo_zs achieves an NMI score of 0.854, demonstrating that the pretrained model can capture regulatory patterns that generalize to a closely related primate species and preserve information relevant for distinguishing macaque cell populations.

Following species-specific post-training, EpiZoo consistently outperforms baseline methods across all four species, covering an evolutionary span from the closely related macaque to the highly divergent zebrafish, fruit fly and maize. Notably, the largest gains are observed in fruit fly and maize, two species for which available scATAC-seq data remain limited in scale and cellular coverage compared with the human and mouse datasets used for pretraining. Despite their greater evolutionary divergence from the pretraining species, EpiZoo improves over the best-performing baselines by 7.0% and 11.2% in fruit fly and maize, respectively. This suggests that EpiZoo does not rely solely on target-species data during post-training, but instead leverages transferable regulatory priors learned from large-scale pretraining. Consistently, UMAP visualizations demonstrate EpiZoo’s improved resolution of fine-grained cell populations that are difficult to distinguish using the baseline methods without large-scale pretraining. In the macaque cortex, for example, the best-performing baseline method, CASTLE, shows limited separation of LAMP5 and LHX6 interneurons, whereas EpiZoo resolves these two closely related subtypes into distinct clusters (Fig. 3b). Similar improvements are observed across evolutionarily divergent species (Fig. 3c-3e). EpiZoo successfully separates the fine-grained cell populations, including Cones, Neurons and Rods in the zebrafish retina, IFM, ISM and TDT muscle subtypes in fruit fly, and Ear and Axillary Bud developmental lineages in maize. Together, these results demonstrate that EpiZoo leverages DNA sequence-derived regulatory priors to adapt across species and generate informative cell representations, particularly when target-species data are limited.

### EpiZoo-Evo decodes conserved and divergent regulatory landscapes across primates

Comparative epigenomic analysis across evolutionary lineages requires resolving how cCREs preserve, diverge or remodel their functions between species. This remains challenging because many distal cCREs, particularly enhancers, show rapid evolutionary turnover in sequence, genomic position and regulatory activity^9,23^, whereas only a subset of cCREs can be reliably connected across species by coordinate mapping. This instability limits the ability of purely coordinate-based methods to identify cCRE-level regulatory conservation and divergence across evolution. Leveraging the DNA sequence-aware architecture of EpiZoo, we developed EpiZoo-Evo to place human and macaque brain epigenomes within a shared pan-primate regulatory space. To support joint analysis of the two species, we constructed a hybrid cCRE vocabulary based on physical genomic overlap after cross-species coordinate mapping, in which cCREs were categorized as shared, human-specific or macaque-specific (Fig. 4a). This vocabulary provided basis for representing conserved and species-specific cCREs together during EpiZoo-Evo post-training. Comprehensive descriptions of the adaptation strategy, including hybrid vocabulary construction, model initialization, and post-training framework for EpiZoo-Evo are provided in Methods.

### EpiZoo-Evo enables joint analysis of cortical cell populations across primates

Raw EpiZoo-Evo cell embeddings reveal separation between human and macaque cortical cells, consistent with global epigenomic divergence accumulated during primate evolution (Fig. 4b). After applying Harmony^47^, a label-free integration method, corresponding cell types from the two species become aligned in the integrated embedding space, while residual separation between species is maintained (Fig. 4c). This pattern indicates that EpiZoo-Evo captures conserved cellular heterogeneity in primates without fully collapsing species-specific epigenomic differences. To further evaluate whether EpiZoo-Evo encode information that can be transferred across species, we performed k-nearest-neighbor label transfer using the raw, unintegrated cell embeddings, as detailed in Methods. Human cells annotated using macaque reference cells achieve an overall accuracy of 0.9095 (Fig. 4d), with an accuracy of 0.9069 even for the rare L6b neuronal subtype, which accounts for only 0.17% of the evaluated cells. Reverse label transfer from human to macaque also achieves an accuracy of 0.8046 (Supplementary Figure 16). Incorrect label assignments are mainly observed among closely related excitatory neuronal lineages. These errors likely arise from the intrinsic similarity of their epigenomic profiles, as well as subtype-specific shifts in chromatin accessibility during species evolution. Together, these results demonstrate that EpiZoo-Evo enables cross-species analysis of cortical cell populations, allowing cell types from closely related species to be compared within a shared embedding space while retaining species-specific epigenomic differences.

**Fig. 4:**
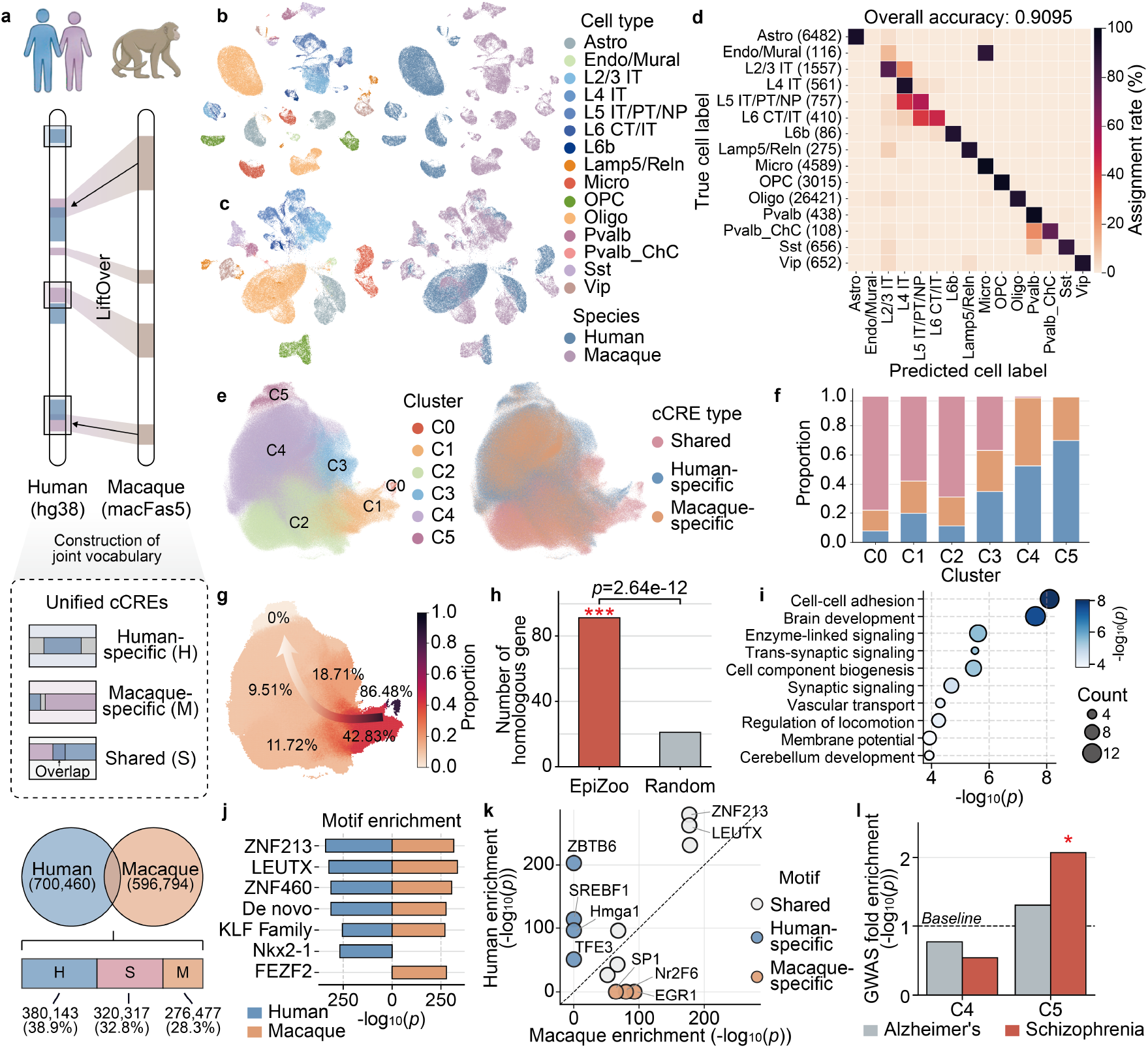
EpiZoo-Evo decodes conserved and divergent regulatory landscapes across primates. **a**, Construction and composition of hybrid cCRE vocabulary. Macaque cCREs are mapped to human genome, and cCREs are categorized as shared, human-specific or macaque-specific according to physical genomic overlap. **b**, UMAP visualizations of raw EpiZoo-Evo cell embeddings labeled by cell type (left) and species (right). **c**, UMAP visualizations of EpiZoo-Evo cell embeddings integrated by Harmony labeled by cell type (left) and species (right). **d**, Cross-species label transfer performance from macaque to human cortical cell types using raw EpiZoo-Evo cell embeddings. Rows indicate true human cell labels, and columns indicate predicted cell labels transferred from macaque reference cells. Numbers in parentheses indicate the number of human cells in each cell type. Color denotes the assignment rate. **e**, EpiZoo-Evo cCRE embeddings projected into a latent manifold and labeled by Louvain cluster or cCRE category. **f**, Composition of shared, human-specific and macaque-specific cCREs across cluster 0 to cluster 5 (denotes as C0 to C5). **g**, Proportion of promoter-proximal shared cCREs across cCRE clusters in the latent space. Promoter-proximal shared cCREs are defined as shared cCREs located within 2.5 kb upstream of a transcription start site in either species. **h**, Homologous gene matching between C4 human-specific cCREs and their nearest macaque-specific neighbors in the EpiZoo-Evo latent space, compared with a random-pairing baseline. Statistical significance was assessed using Fisher’s exact test. Red asterisks denote significant enrichment. **i**, Gene ontology enrichment analysis of homologous genes linked by C4 macro-cCRE pairs. The x-axis and color indicate − log_10_(*p*), and point size indicates gene count. **j**, Motif enrichment in human-specific and macaque-specific cCREs within C4. Direction indicates species-specific enrichment, and bar length indicates − log_10_(*p*). **k**, Motif enrichment comparison between human-specific and macaque-specific cCREs within C5. The x-axis and y-axis indicate − log_10_(*p*) enrichment significance in macaque-specific and human-specific cCREs, respectively. **l**, GWAS enrichment of human-specific cCREs for schizophrenia and Alzheimer’s disease. Statistical significance was assessed using a 1,000-iteration permutation test against the global human cCRE background. Red asterisks denote significant enrichment (*p* < 0.01, permutation test).

### EpiZoo-Evo reveals evolutionary conservation gradients in the cCRE embedding space

Beyond the cell-level relationships described above, regulatory evolution in the primate brain ultimately operates through changes in individual cCREs and their associated functions^11^. We therefore examined the cCRE embeddings learned by EpiZoo-Evo using unsupervised Louvain clustering and quantified the composition of shared, human-specific and macaque-specific cCREs within each cluster. As shown in Fig. 4e and 4f, cCREs are organized into six clusters (cluster 0 to cluster 5, denotes as C0 to C5), which show an overall compositional trend of shared and species-specific cCREs. Across the cCRE embedding space, the proportion of shared cCREs generally decreases from C0 to C5, whereas human-specific and macaque-specific cCREs become more enriched. We further evaluated whether this evolutionary pattern was associated with functional constraint by quantifying the proportion of promoter-proximal shared cCREs within each cluster. These cCREs are expected to be more evolutionarily constrained because of their roles in maintaining core transcriptional programs^48^. The proportion of promoter-proximal shared cCREs decreases from 86.48% in C0 to 0% in C5 (Fig. 4g), following the same trend observed for shared cCRE composition across clusters. These results demonstrate that EpiZoo-Evo cCRE embeddings are biologically interpretable, capturing evolutionary patterns of conservation and divergence among cCREs.

### EpiZoo-Evo identifies putative functional analogies among divergent cCREs across primates

Shared cCREs represent loci that remain mappable across species within conserved genomic context. In contrast, C4 contains almost no shared cCREs, yet human-specific and macaque-specific cCREs are closely co-localized within the EpiZoo-Evo cCRE embedding space. This proximity suggests that a subset of species-specific cCREs may have undergone substantial sequence turnover while retaining related regulatory roles during primate evolution. Inspired by the concept of macro-genes^49^, we refer to these putative functionally analogous cCREs across species as macro-cCREs.

We assessed whether human-specific cCREs and their nearest macaque-specific neighbors in the EpiZoo-Evo latent space were linked to homologous genes, as detailed in Methods. As shown in Fig. 4h, the nearest-neighbor pairing identified 93 human-macaque cCRE pairs whose assigned proximal genes were homologous, significantly exceeding the 21 matches observed under random pairing (*p* = 2.64 × 10^−12^, Fisher’s exact test). Gene ontology enrichment analysis further reveals that these matched genes are associated with conserved nervous system functions, including brain development, cell-cell adhesion and trans-synaptic signaling (Fig. 4i). Because these neurodevelopmental pathways are subject to evolutionary constraint, their associated cCREs may retain similar regulatory roles despite substantial sequence turnover^50^. This linkage between divergent cCREs and conserved biological processes supports the ability of EpiZoo-Evo to identify putatively functionally analogous macro-cCREs.

We further investigated whether these macro-cCREs share convergent regulatory syntax by performing motif enrichment analysis (Fig. 4j). Human-specific and macaque-specific cCREs in C4 are both enriched for motifs from the KLF family, ZNF213, ZNF460, and LEUTX. The enrichment of KLF-family motifs is consistent with reported roles of KLF factors in neuronal maturation^51^, whereas zinc-finger and LEUTX-related paired-like homeobox motifs suggest links to developmental regulatory programs^52,53^. We also observed a shared enriched motif with no clear annotation, suggesting a potentially uncharacterized primate regulatory syntax. Together, the homologous gene mapping and motif enrichment analyses indicate that EpiZoo-Evo can bridge large-scale sequence divergence to recover cross-species regulatory analogies at the cCRE level.

### EpiZoo-Evo identifies divergent regulatory modules associated with human-specific evolution

C5 is separated from the other clusters and contains almost no shared cCREs (Fig. 4e, f), suggesting that it represents a highly divergent regulatory module. Motif enrichment analysis reveals distinct species-specific regulatory signatures within C5 (Fig. 4k). Human-specific cCREs in C5 are enriched for developmental motifs, including ZNF213 and LEUTX, and show human-biased enrichment for Hmga1, SREBF1 and TFE3. Among these human-biased motifs, Hmga1 points to chromatin regulatory programs associated with stemness and progenitor-state regulation^54^, whereas SREBF1 and TFE3 suggest links to lipid metabolism and cellular energetics^55,56^. Together, these motif signatures may connect rapidly evolving human-specific cCREs to chromatin and metabolic regulatory features relevant to cortical development. In contrast, macaque-specific cCREs in C5 are enriched for motifs of broadly acting or context-responsive transcription factors, including SP1, EGR1 and NR2F6. This divergence in motif composition suggests that human-specific cCREs within C5 may represent candidate regulatory programs for recent primate evolutionary innovations.

Evolutionary theory posits that recently evolved regulatory sequences often harbor genetic risk variants for complex human-specific conditions due to the lack of deep ancestral purifying selection^9,57,58^. We evaluated the enrichment of disease risk variants within human-specific cCREs of C4 and C5 using genome-wide association study (GWAS) analysis, as detailed in Methods. As shown in Fig. 4l, human-specific cCREs in C4 show no significant enrichment, consistent with the stronger functional constraint associated with conserved neurodevelopmental regulatory programs. In contrast, human-specific cCREs in C5 are significantly enriched for schizophrenia-associated variants (*p* < 0.01, permutation test), but not for the Alzheimer’s disease-associated variants. This selective enrichment links C5 to recently evolved, human-specific regulatory programs rather than broadly conserved neurological pathways, consistent with the stronger evolutionary and neurodevelopmental component of schizophrenia risk compared with Alzheimer’s disease. Together, these analyses demonstrate that EpiZoo-Evo can resolve regulatory conservation, divergence and functional analogy across primate epigenomes.

### EpiZoo-Cancer prioritizes somatic mutations in cancer-type-specific regulatory contexts

Single-nucleotide variants in noncoding genomic regions frequently disrupt transcription factor binding, alter chromatin accessibility and contribute to oncogenesis. Although existing single-cell foundation models effectively capture cellular heterogeneities, they generally lack explicit DNA sequence encoding, limiting their ability to evaluate the regulatory effects of nucleotide-level perturbations. EpiZoo addresses this limitation through SEAM, which encodes the underlying DNA sequence of each cCRE and enables the prediction of how single-nucleotide variants affect chromatin accessibility. We extended EpiZoo to oncology by developing EpiZoo-Cancer, a context-aware framework that prioritizes noncoding somatic mutations that overlap cCREs using loss-of-accessibility (LoA) scores (Fig. 5a). In this framework, each cancer type is represented by a learnable cancer-type embedding, which allows EpiZoo to capture cancer-type-specific regulatory programs during fine-tuning on a pan-cancer scATAC-seq dataset^59^. The detailed datasets, fine-tuning strategy and LoA computation are described in Methods.

We computed LoA scores for 5,959 variants overlapping human cCREs and mapped each variant to its nearest gene, as detailed in Methods; the complete set of evaluated variants is provided in Supplementary Table 4. We then assessed whether the high-impact mutations prioritized by EpiZoo-Cancer were mapped to genes defined as Tier 1 cancer driver genes in the Cancer Gene Census (CGC) from the Catalogue of Somatic Mutations in Cancer (COSMIC), a curated resource of cancer driver genes^60^. As shown in Fig. 5b and 5c, high-impact mutations prioritized by both zero-shot and fine-tuned EpiZoo-Cancer are consistently enriched for assignments to known cancer driver genes. For instance, in breast invasive carcinoma (BRCA), zero-shot and fine-tuned EpiZoo-Cancer show significant enrichment (*p* = 8.59 × 10^−3^ and *p* = 4.69 × 10^−3^, Fisher’s exact test), with high-impact mutations assigned to 19 and 20 cancer-related genes, respectively. Similarly, in kidney renal clear cell carcinoma (KIRC), both models show enrichment for variants assigned to driver genes, with odds ratios of 1.75 and 2.26. These results indicate that EpiZoo-Cancer can prioritize candidate regulatory disruptions across diverse cancer types through explicit DNA sequence modeling. The strong zero-shot performance further suggests that pretraining on normal epigenomic profiles allows the model to learn generalizable patterns of chromatin accessibility regulation and identify somatic variants that perturb these regulatory programs without cancer-type-specific fine-tuning.

**Fig. 5:**
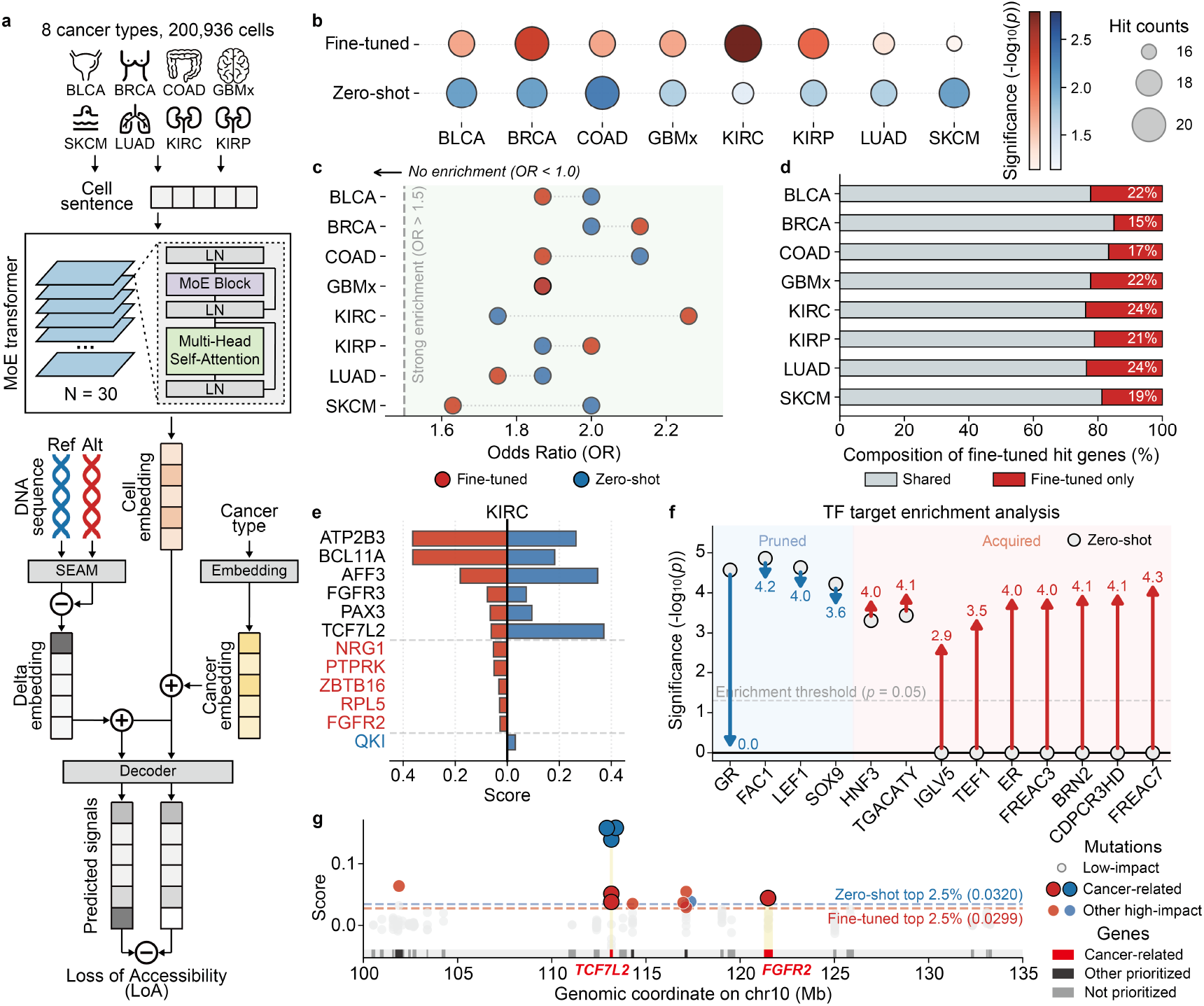
EpiZoo-Cancer prioritizes somatic mutations in cancer-type-specific regulatory contexts. **a**, Overview of the EpiZoo-Cancer framework. A learnable cancer-type embedding is integrated with cell embeddings to provide cancer-type-specific regulatory context for computing LoA scores. Cancer abbreviations: BLCA, bladder urothelial carcinoma; BRCA, breast invasive carcinoma; COAD, colon adenocarcinoma; GBMx, glioblastoma multiforme; SKCM, skin cutaneous melanoma; LUAD, lung adenocarcinoma; KIRC, kidney renal clear cell carcinoma; KIRP, kidney renal papillary cell carcinoma. LN, layer normalization. **b**, Enrichment of high-impact mutations assigned to COSMIC CGC Tier 1 driver genes across eight cancer types. Point size indicates the number of driver-gene hits, and color denotes enrichment significance from Fisher’s exact test. **c**, Odds ratios for the enrichment analysis in **b. d**, Candidate genes prioritized by the fine-tuned EpiZoo-Cancer, including genes prioritized exclusively after fine-tuning and genes shared with zero-shot EpiZoo-Cancer. **e**, LoA scores of high-impact KIRC mutations assigned to cancer-related genes. Values on the left indicate predictions from fine-tuned EpiZoo-Cancer, and values on the right indicate predictions from zero-shot EpiZoo-Cancer. **f**, Differential transcription factor target enrichment analysis for KIRC variants prioritized by zero-shot and fine-tuned EpiZoo-Cancer. Arrows indicate changes in enrichment significance after cancer-type-specific fine-tuning, and the dashed line indicates the significance threshold. **g**, Distribution of LoA scores on chromosome 10. Each point represents an individual variant, and the dashed line indicates the threshold used to define high-impact mutations.

Although the zero-shot EpiZoo-Cancer captures regulatory variants with broad effects, tumorigenesis is shaped by distinct regulatory contexts across cancer types. By integrating cancer-type embeddings, fine-tuned EpiZoo-Cancer adjusts variant prioritization and identifies additional candidate disrupted genes. Across the eight evaluated cancer types, the fine-tuned model prioritizes an average of 20.5% additional genes that are not detected by zero-shot EpiZoo-Cancer (Fig. 5d). In KIRC, both models identify high-impact mutations assigned to *TCF7L2* and *FGFR3* (Fig. 5e). These genes are associated with Wnt/β-catenin and fibroblast growth factor (FGF)-fibroblast growth factor receptor (FGFR) signaling pathways, respectively^61,62^. The fine-tuned model uniquely prioritizes *PTPRK* and *FGFR2*, which are associated with epithelial adhesion, tumor-suppressive signaling and growth factor receptor pathways, suggesting that cancer-type-specific fine-tuning captures regulatory effects that better reflect the biological context of each cancer type^63,64^. Conversely, *QKI*, identified only by the zero-shot model, encodes an RNA-binding protein involved in broad post-transcriptional regulation, suggesting that zero-shot predictions may preferentially capture more general regulatory disruptions^65^.

We next evaluated transcription factor enrichment signatures associated with KIRC mutations prioritized by EpiZoo-Cancer (Fig. 5f). Compared with zero-shot EpiZoo-Cancer, the fine-tuned model shows reduced enrichment for broadly acting regulators such as the glucocorticoid receptor (GR) and increased enrichment for factors associated with disease including TEF1 from the TEAD family^66^ and HNF3 from the FOXA family^67^. GR is broadly involved in glucocorticoid-mediated stress-response programs^68^, whereas TEAD family and FOXA family target programs point to Hippo-YAP/TAZ signaling and epithelial lineage regulation^66,69^. This shift suggests that the fine-tuning strategy enables EpiZoo-Cancer to better capture regulatory patterns associated with different cancer types. Mapping LoA scores along genomic coordinates further shows that high-impact mutations are assigned to validated cancer driver genes, consistent with the enrichment analysis above (Fig. 5g). Collectively, EpiZoo-Cancer integrates DNA sequence syntax with disease context to prioritize candidate cancer-related regulatory disruptions, extending EpiZoo from single-cell epigenomic feature extraction to nucleotide-level interpretation of somatic variants.

### EpiZoo translates DNA sequence syntax into cell-type-specific chromatin accessibility

Most sequence-based models predict regulatory activity from DNA sequence alone or under predefined cellular contexts, limiting their ability to incorporate cell-type-specific regulatory information. Benefiting from its ability to jointly encode DNA sequence-derived regulatory information and cellular heterogeneity, EpiZoo can be extended to predict cell-type-specific chromatin accessibility by combining SEAM-derived sequence embeddings with cell-type embeddings, which were obtained by aggregating cell embeddings within each cell type (Fig. 6a). We used the mouse brain (BICCN2) dataset provided by the CREsted framework^70^, which categorized scATAC-seq data across 19 cell types into 49,936 consensus peaks and 8,198 cell-type-specific peaks, respectively. The detailed framework design and training strategy are provided in Methods.

**Fig. 6.**
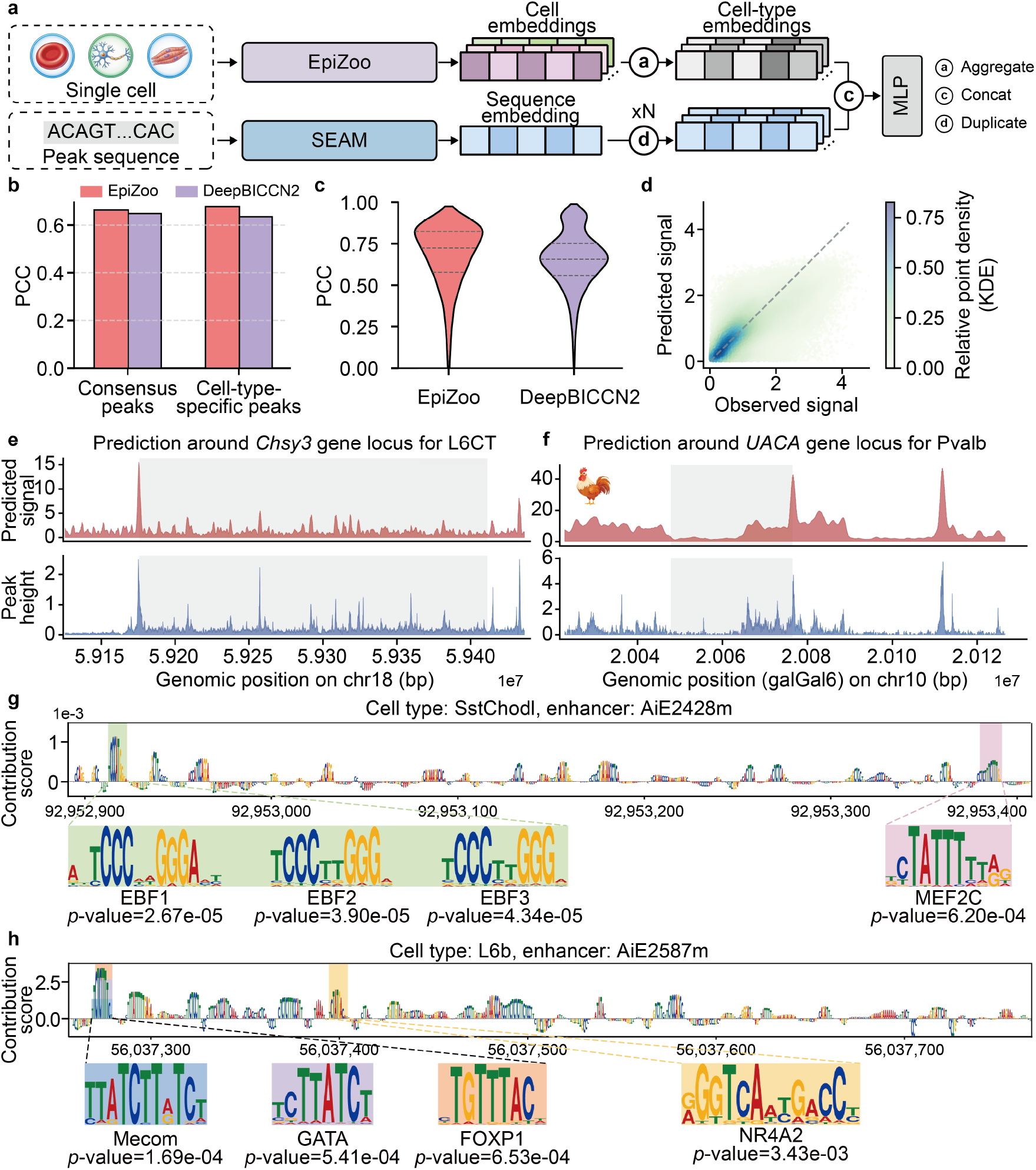
EpiZoo translates DNA sequence syntax into cell-type-specific chromatin accessibility. **a**, Overview of the cell-type-specific chromatin accessibility prediction framework. Cell-type embeddings derived from EpiZoo are concatenated with sequence embeddings generated by SEAM to predict chromatin accessibility across different cell types. **b**, Cell-type-level PCCs between predicted and observed chromatin accessibility signals for EpiZoo and DeepBICCN2 on consensus and cell-type-specific peaks. **c**, Peak-level PCC distributions for cell-type-specific peaks. The inner horizontal lines denote the first quartile, median, and third quartile. **d**, Relative point density of predicted and observed chromatin accessibility signals measured by kernel density estimation (KDE). **e**, Continuous chromatin accessibility prediction across the mouse *Chsy3* gene locus in L6 corticothalamic (L6CT) neurons. **f**, Cross-species chromatin accessibility prediction across the chicken *UACA* gene locus using the mouse Pvalb interneuron cell type context. **g**, Nucleotide-resolution gradient attribution for the AiE2428m enhancer in SstChodl interneurons, with high-attribution regions annotated by candidate transcription factor binding motifs. **h**, Nucleotide-resolution gradient attribution for the AiE2587m enhancer in L6b projection neurons, with high-attribution regions annotated by candidate transcription factor binding motifs.

We benchmarked EpiZoo on held-out test peaks against DeepBICCN2, the baseline model established by CREsted^70^, measured by PCCs (Fig. 6b). At the cell-type level, EpiZoo achieves PCCs of 0.663 and 0.677 on consensus and cell-type-specific peaks, respectively, outperforming the corresponding baseline scores of 0.648 and 0.634. Peak-level analysis further demonstrates that EpiZoo more accurately captures quantitative differences in chromatin accessibility across cell types, increasing the median PCC from 0.646 to 0.706 (Fig. 6c). These improvements indicate that the explicit introduction of cell-type embeddings enables EpiZoo to decode how the same DNA sequence can exhibit distinct accessibility signals across different cell types. Consistently, the predicted and observed signals show strong alignment along the diagonal in the density plot, further supporting the quantitative fidelity of the EpiZoo’s prediction (Fig. 6d).

We then examined whether this prediction could extend beyond predefined peaks to reconstruct continuous chromatin accessibility landscapes. Using a sliding-window inference strategy, EpiZoo sequentially predicts the chromatin accessibility signals across genomic segments. At the mouse *Chsy3* locus in L6 corticothalamic (L6CT) neurons, the predicted signals closely matched the observed peak heights, with a PCC of 0.647 (Fig. 6e). Furthermore, we evaluated cross-species sequence interpretation by predicting the chromatin accessibility across the chicken *UACA* gene locus. The sequence embeddings extracted from the galGal6 genome are concatenated with the mouse Pvalb interneuron cell-type embedding. Our model reconstructs an accessibility profile concordant with the observed signals, with a PCC of 0.657, suggesting that sequence-derived regulatory features can be interpreted under a transferred mammalian neuronal context (Fig. 6f).

Next, we performed nucleotide-resolution gradient attribution to identify motifs contributing to cell-type-specific enhancer activity, as detailed in Methods. For the AiE2428m enhancer in SstChodl interneurons, high-attribution regions overlap candidate EBF-family and MEF2C binding motifs, transcription factors with established roles in neurogenesis and cortical neuronal development^71,72^ (Fig. 6g). For the AiE2587m enhancer in L6b projection neurons, high-attribution regions overlap candidate FOXP1, Mecom and GATA-family motifs (Fig. 6h). Because L6b projection neurons are closely related to the transient embryonic subplate^73^, the recovery of these developmental motifs suggests that EpiZoo captures sequence features associated with lineage history and developmental context.

Collectively, by combining SEAM-derived sequence embeddings with learned cell-type embeddings, EpiZoo links nucleotide-level regulatory grammar to epigenomic activity in specific cellular contexts and supports regulatory interpretation across cell types, genomic regions and species.

## Discussion

In this study, we introduce EpiZoo, a DNA sequence-aware foundation model for cross-species single-cell epigenomics. Pretrained on the Omni-scATAC corpus, EpiZoo learns generalizable regulatory landscapes from large-scale human and mouse scATAC-seq profiles. By combining DNA sequence-derived regulatory information, sequence-independent epigenomic context and accessibility-based importance, EpiZoo captures cellular heterogeneity across diverse datasets while reducing dependence on fixed species-specific genomic coordinates. The capability supports robust feature extraction, cell type annotation and data imputation, demonstrating the utility of EpiZoo as a general framework for fundamental single-cell analysis.

The main advance of EpiZoo is to incorporate DNA sequence information and cross-species framework into foundation modeling of single-cell chromatin accessibility. SEAM-derived sequence embeddings provide transferable regulatory priors for cCREs, while multi-species pretraining enables EpiZoo to learn both conserved regulatory principles and species-specific chromatin accessibility patterns. This design allows EpiZoo to adapt to evolutionarily diverse species and extends single-cell epigenomic foundation modeling beyond cell representation learning toward sequence regulatory interpretation. Through this framework, EpiZoo supports comparative analysis of regulatory evolution, context-aware prioritization of somatic variants and prediction of cell-type-specific chromatin accessibility from DNA sequences.

We outline three future directions to extend EpiZoo. First, EpiZoo can be expanded from single-cell epigenomics to multi-omics regulatory modeling. By incorporating transcriptomic, proteomic and other phenotypic measurements, future models could learn how chromatin accessibility, gene expression and protein abundance are coordinated across tissues, species and biological contexts. Second, the DNA sequence-aware design of EpiZoo provides a foundation for systematic interpretation of regulatory variants. Extending this framework to larger variant catalogs, disease cohorts and experimentally validated perturbation datasets would help assess how noncoding sequence changes alter chromatin accessibility and downstream molecular programs. Third, EpiZoo can be further developed into a predictive framework for regulatory perturbation analysis. By integrating datasets involving genetic perturbations and pharmacological treatments, future models could predict how regulatory programs respond under specific cellular contexts. This would extend EpiZoo from representation and interpretation toward predictive modeling of disease mechanisms and therapeutic interventions, providing a foundation for virtual cell frameworks.

## Methods

### Omni-scATAC: a comprehensive scATAC-seq corpus for pretraining

We assembled Omni-scATAC as the data foundation for EpiZoo by manually curating available human and mouse scATAC-seq datasets from published studies. The human component included the pretraining datasets used by EpiAgent^20^, which comprise approximately 4.8 million cells and constitute the major part of the Human-scATAC-Corpus^74^, together with 12 additional human datasets curated in this study^28,29,75-84^. These additional datasets expanded the human component to approximately 8.1 million cells. For mouse, we collected 33 public datasets spanning diverse biological contexts and experimental protocols^13,30,77,85-113^, comprising approximately 12.8 million cells. In total, Omni-scATAC comprised 46 dataset entries and approximately 20.9 million scATAC-seq profiles spanning more than 30 tissues or cell lines. We organized data availability, source species, tissue or cell line, sequencing technology and data statistics with detailed information provided in Supplementary Table 1.

The scATAC-seq data were converted into standardized cell-by-cCRE matrices within species-specific feature spaces. Human cCREs were based on the vocabulary established by Human-scATAC-Corpus, whereas the mouse cCRE vocabulary was curated from a large-scale mouse scATAC-seq atlas^96^ in the pretraining data. The coverage of the human cCRE vocabulary has been systematically evaluated in EpiAgent and Human-scATAC-Corpus, where overlap analyses showed that the curated human cCREs cover the large majority of independently collected human peak and cCRE sets. We performed the same assessment for the mouse cCRE vocabulary against independent mouse peak and cCRE collections, and observed similarly broad coverage. Before model training, we further applied cCRE filtering for both human and mouse cCREs. Detailed cCRE construction, filtering and assessment are provided in Supplementary Note 3 and Supplementary Figure 17.

### Cell sentence construction

EpiZoo represented each cell as a cell sentence. Human and mouse cells were processed separately using their corresponding species-specific cCRE vocabularies. For each cell, accessible cCREs were ranked in descending order according to the global TF-IDF scores computed with a precomputed document-frequency vector from the corresponding subset of pretraining corpus. The cell sentence began with a [CLS] token, followed by the ranked accessible cCRE tokens, and ended with a [SEP] token. The maximum input length of EpiZoo was set to *L* = 8192. This setting follows EpiAgent^20^, which showed that a maximum token length of 8,192 provides an effective balance between cCRE coverage and computational efficiency, with limited additional gains from longer inputs. If the number of accessible cCREs exceeded *L*− 2, with two positions were reserved for [CLS] and [SEP], accessible cCREs were randomly sampled and reordered according to the original TF-IDF ranking. Detailed global TF-IDF calculation is provided in Supplementary Note 4.

For each cell, the cell sentence is defined as:

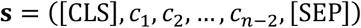

where *n* denotes the length of the unpadded cell sentence, and *c*_*t*_ denotes the *t*-th cCRE token ordered by the global TF-IDF scores. Because cCRE vocabularies are species-specific, *c*_*t*_ is drawn from the human cCRE vocabulary *V*^h^ for human cells and from the mouse cCRE vocabulary *V*^m^ for mouse cells.

### EpiZoo model architecture

The overall architecture of EpiZoo contains approximately 2.6 billion parameters in total and comprises three core modules: a DNA sequence-aware embedding module, a MoE transformer and species-specific signal decoders. The DNA sequence-aware embedding module transformed the input cell sentence into a series of token embeddings. The MoE transformer then contextualized these token embeddings across the cell sentence, and the output vector corresponding to the [CLS] token was used as the cell embedding. The species-specific signal decoders mapped this cell embedding to the corresponding cCRE feature space.

### DNA sequence-aware embedding module

The module mapped each token into an embedding vector by combining three components: a sequence embedding, a learnable identity embedding and a learnable rank embedding. The sequence embedding was generated by SEAM, which consisted of a DNABERT-2^114^ model followed by a linear projection head. For each cCRE *c*, SEAM encoded its DNA sequence to derive the sequence embedding:

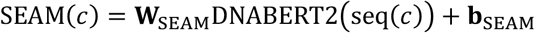

where seq(*c*) denotes the DNA sequence of cCRE *c* within its corresponding genome, DNABERT2(·) denotes the DNABERT-2 model, W_SEAM_ and b_SEAM_ denote parameters of SEAM’s projection head. For special tokens, including [CLS], [SEP] and [PAD], the placeholder sequence embedding was represented by vectors inherited from its corresponding cCRE identity embedding. After multi-round calibration that iteratively aligned SEAM-derived sequence embeddings with the EpiZoo cCRE embedding space, SEAM was frozen for subsequent procedures. For standard downstream fine-tuning and inference on the pretraining vocabularies, the sequence embeddings were precomputed and used directly. For species-specific or cross-species post-training involving newly defined cCRE vocabularies, the frozen SEAM was applied to the DNA sequences of the new cCREs to generate the corresponding sequence embeddings. These newly computed sequence embeddings were then fixed and used as the sequence-derived component of the token embeddings during post-training.

For each retained cCRE token *c*_*t*_, the token embedding was formed by summing three components: the SEAM-derived sequence embedding, the identity embedding and the rank embedding:

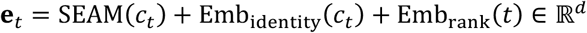

where *d* = 512 denotes the hidden size of EpiZoo, *c*_*t*_ denotes the *t*-th cCRE token ordered by the global TF-IDF scores, *t* denotes its rank position in the cell sentence, and Emb_identity_(·) and Emb_rank_(·) denote the lookup functions of the learnable identity embedding matrix and rank embedding matrix, respectively. The token embeddings generated by the DNA sequence-aware embedding module were stacked in the order of the cell sentence to form the input embedding matrix for the MoE transformer:

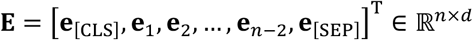

where *n* = 8192 denotes the input length and *d* = 512 denotes the hidden size.

### MoE transformer

We set the token embedding matrix E as the initial hidden representation of the MoE transformer, denoted as **H**^(0)^. In each transformer block *l*, the hidden representation was first updated by the multi-head self-attention layer, yielding an intermediate representation 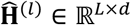:

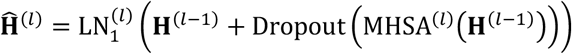

where LN(·) denotes layer normalization, MHSA(·) denotes multi-head self-attention computation. MoE layer was then applied to the intermediate representation matrix:

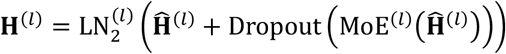

where MoE(·) denotes the computation of the MoE layer, which was applied independently to each token representation. Specifically, for each token representation h ∈ ℝ^*d*^ from 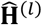, the gating network computed routing probabilities over *N*_exp_ = 4 experts:

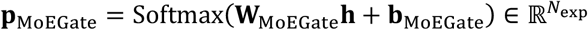

where **W**_MoEGate_ and **b**_MoEGate_ denote parameters of the MoE gating network. Let *E* denote the set of top-*K* selected experts based on the routing probabilities, where *K* = 2. The routing probabilities of the selected experts were renormalized as:

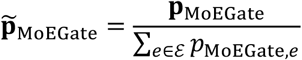

The output of MoE layer was computed as the weighted sum of the selected expert outputs:

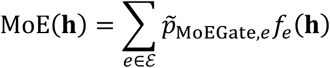

where *f*_*e*_(·) denote the feed-forward network of expert *e*, implemented as a two-layer network with GELU activation and dropout. This operation was applied independently to all token representations in the hidden representation matrix. The outputs were then stacked back in the original order to form the MoE output matrix, which was used to update the hidden representation and served as the input to the next block. After the final block, the [CLS] token embedding was extracted as the cell embedding:

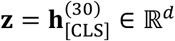

### Species-specific signal decoder

The species-specific signal decoders reconstructed accessibility signals from the cell embedding within the corresponding cCRE vocabulary. The decoder was implemented as a collection of independent branches, with one branch assigned to each species:

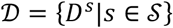

where *S* denotes the set of species and *D*^*s*^ denotes the species-specific signal decoder. For a cell from species *s*, only the corresponding decoder branch was activated:

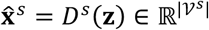

where 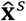 denotes the reconstructed signals and k^*s*^ denotes the cCRE vocabulary of species *s*. Each species-specific signal decoder was implemented as a linear projection:

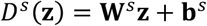

where 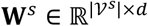 and 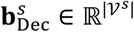 denote the parameters of the species-specific signal decoder.

During pretraining and representation-learning tasks, the species-specific decoder was trained to reconstruct the binary accessibility profile within the matched cCRE vocabulary, with the decoder output used as logits for binary reconstruction loss. In data imputation, the same decoder branch was fine-tuned against TF-IDF signals using mean squared error loss, so that the reconstructed output recovered continuous accessibility values.

### Pretraining strategy

EpiZoo was pretrained using a DNA sequence-aware multi-stage strategy that couples SEAM calibration with species-specific self-supervised learning. We first calibrated SEAM to map cCRE DNA sequences into the latent embedding space of EpiZoo. The main model was then optimized using two self-supervised tasks, including cell-cCRE alignment and signal reconstruction. These tasks were adapted from the EpiAgent pretraining framework^20^, in which ablation analysis have demonstrated the individual and combined contributions of these tasks to the model performance. In EpiZoo, both tasks were performed within matched species-specific cCRE vocabularies, so that cells were aligned only with cCREs from the same species and reconstructed only through the corresponding species-specific signal decoder branch.

#### Cell-cCRE alignments

The cell-cCRE alignment task trained the cell embedding to predict the accessibility of sampled cCREs from the same species. For each cell from species *s*, let 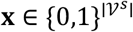 denote the binary accessibility vector over the species-specific cCRE vocabulary k^*s*^. The accessible and inaccessible cCRE sets were defined as:

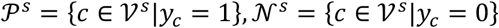

We then sampled a balanced set of positive and negative cCREs:

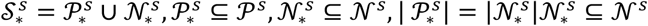

For each sampled 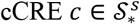, the cell embedding and the cCRE identity embedding were concatenated:

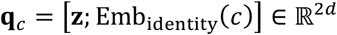

where [·;·] denotes vector concatenation. The concatenated representation was passed to the cell-cCRE alignment head to obtain the accessibility logit:

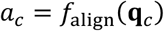

where *f*_align_(·) was implemented as a two-layer multilayer perceptron with GELU activation, layer normalization and dropout. The alignment task was optimized using binary cross-entropy loss:

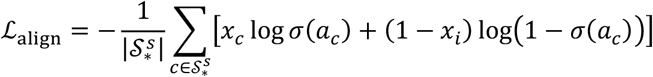

where *σ*(·) denotes the sigmoid function.

#### Species-specific signal reconstruction

The signal reconstruction task trained the cell embedding to reconstruct the binary accessibility signals in the corresponding species-specific cCRE vocabulary. For each cell from species *s*, the reconstruction loss was computed between the output of signal decoder branch 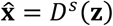 and the binary accessibility vector **x**:

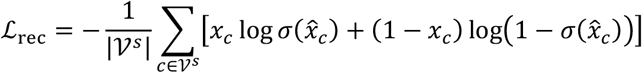

The final pretraining loss function was the sum of these two losses:

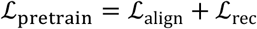

#### SEAM calibratio

During the initial training epochs, EpiZoo was trained without introducing the SEAM-derived sequence embedding component, so that each cCRE token was represented only by its learnable identity embedding and rank embedding. The identity embeddings obtained from this initial training stage were then used as regression targets to calibrate SEAM. For cCRE *c* from species *s*, let Emb_identity_(*c*) denote the cCRE identity embedding learned during this stage. SEAM was optimized using mean squared error loss:

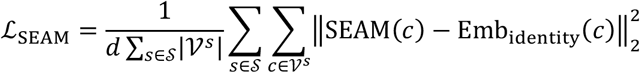

After this calibration, SEAM and the remaining EpiZoo parameters were alternately frozen and updated in the initial stage of pretraining under the cell-cCRE alignment and signal reconstruction tasks described above. SEAM was then frozen during the subsequent pretraining, downstream fine-tuning and inference stages, except for the cell-type-specific chromatin accessibility prediction.

### Fundamental downstream tasks for single-cell analysis

We evaluated EpiZoo on three fundamental downstream tasks for single-cell chromatin accessibility analysis, including feature extraction, cell type annotation and data imputation.

#### Feature extraction

For feature extraction, EpiZoo was fine-tuned on each benchmark dataset using the same tasks as pretraining, including cell-cCRE alignment and species-specific signal reconstruction. We then evaluated fine-tuning strategies across the three core modules of EpiZoo. Each module was assigned either FFT or LoRA, yielding eight configurations. Detailed LoRA formulations and ablation settings are provided in Supplementary Note 5. For clustering evaluation, Louvain clustering was performed on the cell embeddings. Clusters were compared with reference cell labels using NMI and ARI, with formulations provided in Supplementary Note 6. For trajectory analysis, EpiZoo embeddings of B-lineage cells from Cusanovich2018 were used to construct the neighborhood graph, followed by PAGA-based trajectory inference and DPT analysis^35^.

#### Cell type annotation

For cell type annotation, a classifier head was appended to the EpiZoo cell embedding. The classifier head was implemented as a three-layer multilayer perceptron and produced predicted probabilities over cell types. EpiZoo and the classifier head were jointly fine-tuned using focal loss with ground-truth cell type labels.

The annotation task was evaluated under intra-dataset and inter-dataset settings. For intra-dataset validation, each dataset was divided into five folds, with four folds used for training and one fold used for testing. For inter-dataset validation, EpiZoo was fine-tuned on reference data from the same tissue context in the Omni-scATAC corpus and tested on external datasets. Specifically, Kanemaru2023 was tested after training on the cardiac subset of Zhang2021; Li2023b after training on Li2023a; Cusanovich2018 after training on Lu2025 and Zu2023; and Fang2021 after training on Zu2023. Performance was evaluated using accuracy, κ and mF1 score.

#### Data imputation

For data imputation, we standardized the cell-by-cCRE matrices to ensure a fair comparison. Following the parameter setting used in scCASE, we retained cCREs accessible in at least 3% of cells for each benchmark dataset. The same filtering threshold was applied to EpiZoo and all baseline methods. The retained accessibility matrix was then transformed into TF-IDF normalized signals and used as the reference matrix.

To simulate dropout, 50% of the nonzero entries in the reference matrix were randomly masked to generate a corrupted input matrix. EpiZoo was fine-tuned to reconstruct the original TF-IDF normalized accessibility signals from the corrupted input using the species-specific signal decoders, and the reconstructed output was used as the imputed accessibility matrix. Baseline methods were run using the parameter settings described in their original studies or their default implementations. The imputation task was evaluated by Louvain clustering on the imputed matrices and by PCCs between imputed and reference signals at the cell type, cell and cCRE levels. Detailed clustering metrics, classification metrics and PCC calculations are provided in Supplementary Note 6.

### Species-specific adaptation of EpiZoo to new species

We adapted the pretrained EpiZoo to new species through three steps, consisting of sequence embedding computation, model initialization and post-training. First, SEAM was applied to the DNA sequences of cCREs in the new species to compute sequence embeddings. Second, EpiZoo was initialized by transferring available pretrained parameters according to evolutionary distance. Third, the initialized model was post-trained on scATAC-seq data from the new species. In this study, we demonstrate this adaptation using macaque, zebrafish, fruit fly and maize as representative examples spanning different evolutionary distances.

#### Sequence embedding computation

For a target species *s*, let *V*^*s*^ denote its cCRE vocabulary. For each cCRE *c* ∈ *V*^*s*^, its DNA sequence was extracted from the corresponding genome and encoded by SEAM to generate sequence embedding. The resulting sequence embeddings were used as the sequence-derived component of token embeddings for the new species. During species adaptation, SEAM remained frozen.

#### Model initialization

For phylogenetically close species, cCREs of the target species were mapped to a reference species using UCSC liftOver^115^. Let *s* denote the new species, *r* denote the reference species, *V*^*s*^ and *V*^*r*^ denote their cCRE vocabularies. We defined a coordinate mapping

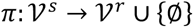

where *π*(*c*) = *c*^′^ indicates that cCRE *c* from the new species overlaps reference cCRE *c*^′^ after liftOver, and *π*(*c*) = ∅ indicates that no reference cCRE was matched.

For each cCRE *c* ∈ k^*s*^, its identity embedding was initialized as:

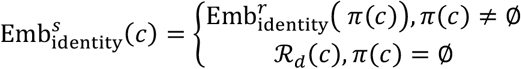

where 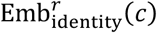 denotes the pretrained cCRE identity embeddings of the reference species, and *R*_*d*_(·) denotes a random initialization function that returns a *d*-dimensional embedding vector. For the signal decoder *D*^*s*^, parameters corresponding to mapped cCREs were inherited from the matched reference cCREs, whereas parameters corresponding to unmapped cCREs were randomly initialized. This strategy allowed EpiZoo to reuse the regulatory information learned from pretraining while accommodating newly defined cCREs in the new species.

For evolutionarily distant species without reliable liftOver to the pretraining species, including zebrafish, fruit fly and maize, inheritance based on coordinates was not used. In these cases, all cCRE identity embeddings and decoder parameters for the new specie were randomly initialized. Parameters shared across species, including the MoE transformer, rank embeddings and cell-cCRE alignment head, were inherited from the pretrained EpiZoo model. For the zero-shot macaque evaluation, no parameter update was performed; cell embeddings were generated using only macaque cCREs that could be mapped to human cCREs, together with the inherited pretrained parameters.

#### Post-training

After model initialization, EpiZoo was post-trained on scATAC-seq data from the target species using same tasks as pretraining, including cell-cCRE alignment and signal reconstruction. Both tasks were performed within the cCRE vocabulary of the corresponding species. After post-training, the cell embedding for each cell was extracted from the output vector of the [CLS] token in the MoE transformer. Embedding quality was evaluated by Louvain clustering, followed by comparison with reference cell type annotations using NMI and ARI. UMAP was used for visualization^116^.

### EpiZoo-Evo for modeling primate regulatory evolution

EpiZoo-Evo was developed to jointly model human and macaque cortical scATAC-seq data within a shared primate cCRE vocabulary. This workflow consisted of hybrid vocabulary construction, model initialization, post-training, and analysis of the resulting cell and cCRE embeddings.

#### Hybrid vocabulary construction

Let *V*^h^ and *V*^m^ denote the human and macaque cCRE vocabularies, respectively. Macaque cCREs were mapped to the human hg38 genome using UCSC liftOver. We defined the mapping from macaque to human as:

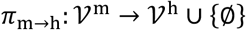

where *π*_m→h_(*c*^m^) = *c*^h^ indicates that macaque cCRE *c*^m^ overlapped human cCRE *c*^h^ after liftOver, and *π*_m→h_(*c*^m^) = ∅ indicates that no overlapping human cCRE was found.

Human cCREs with at least one mapped macaque cCRE were defined as shared cCREs:

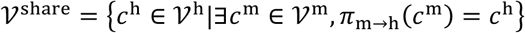

The remaining human cCREs were defined as human-specific cCREs, and macaque cCREs without mapped human cCREs were defined as macaque-specific cCREs:

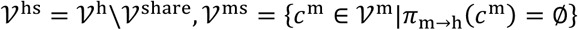

The EpiZoo-Evo hybrid vocabulary was defined as *V*^evo^ = *V*^share^ ∪ *V*^hs^ ∪ *V*^ms^.

#### Model initialization

For each cCRE in the hybrid vocabulary *V*^evo^, its DNA sequence was extracted from the corresponding genome and encoded by SEAM to generate sequence embedding. The cCRE identity embeddings in EpiZoo-Evo were initialized according to cCRE category:

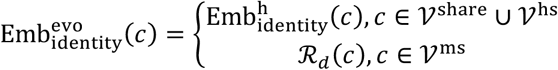

where 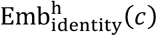 denotes the pretrained human cCRE identity embedding, and *R*_*d*_(·)denotes a random initialization function that returns a *d*-dimensional embedding vector. The EpiZoo-Evo signal decoder was defined over the hybrid vocabulary. Decoder parameters corresponding to shared and human-specific cCREs were inherited from the pretrained human signal decoder branch, whereas parameters corresponding to macaque-specific cCREs were randomly initialized. Parameters shared across species, including the MoE transformer, rank embeddings and cell-cCRE alignment head, were inherited from pretrained EpiZoo.

#### Post-training

EpiZoo-Evo was post-trained on the integrated human and macaque cortex scATAC-seq dataset using the same self-supervised tasks as pretraining, with both losses computed within the hybrid vocabulary. After post-training, the cell embedding for each cell was extracted from the output vector of the [CLS] token in the MoE transformer. For visualization of cross-species cell embeddings, raw EpiZoo-Evo cell embeddings were projected using UMAP.

#### Analysis of EpiZoo-Evo cell embeddings

Harmony integration was applied to raw cell embeddings using species as the batch covariate for joint visualization. Label transfer was performed using raw, unintegrated EpiZoo-Evo cell embeddings in both human-to-macaque and macaque-to-human directions. Cell embeddings from each species were independently mean-centered and L2-normalized before nearest-neighbor search. For each query cell, the top *k* nearest reference cells were identified using cosine similarity, and the query label was assigned by majority voting among the corresponding reference labels. Detailed harmony integration, cross-species label transfer strategy and assessment are provided in Supplementary Note 7.

#### Analysis of EpiZoo-Evo cCRE embeddings

For cCRE analyses, we used the cCRE identity embeddings learned by EpiZoo-Evo after post-training. These embeddings were projected using UMAP and clustered using Louvain clustering. Promoter-proximal cCREs were identified by genomic intersection using pybedtools.

For each shared cCRE *c* ∈ *V*^share^, the corresponding macaque cCREs were defined according to the mapping from macaque to human:

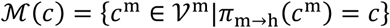

For each species, promoter regions were defined as the 2.5 kb upstream regions of annotated transcription start sites. A shared cCRE was defined as promoter-proximal if the midpoint of its human genomic interval overlapped a human promoter region, or if the midpoint of any corresponding macaque cCRE in *M*(*c*) overlapped a macaque promoter region.

Homologous gene mapping was performed for human-specific and macaque-specific cCREs within selected EpiZoo-Evo cCRE clusters. For each human-specific cCRE, cosine similarity to macaque-specific cCREs was computed using the learned cCRE embeddings, and top *k* nearest macaque-specific cCREs were selected to construct cross-species cCRE pairs (*k* = 5). Each cCRE was assigned to its nearest proximal gene in the corresponding species. A cCRE pair was considered homologous if the assigned human and macaque genes were annotated as homologous genes. Homologous gene annotations between human and macaque were obtained from Ensembl BioMart^117^. Randomly paired human-specific and macaque-specific cCREs were used as the background, and statistical significance was assessed using a one-sided Fisher’s exact test, as detailed in Supplementary Note 8.

Gene ontology enrichment analysis was performed using Metascape^118^. Homologous genes identified from C4 macro-cCRE pairs were used as the input gene list. Functional enrichment was performed against Gene Ontology biological process terms using the default Metascape background. Enriched terms were ranked according to their enrichment values reported by Metascape, and representative biological process terms were selected for visualization.

Motif enrichment analysis was performed using XSTREME^119^ from the MEME Suite online server^120^. For each cCRE group, including human-specific and macaque-specific cCREs within C4 or C5, genomic sequences were extracted and submitted as the primary sequence set. XSTREME was used for motif discovery, enrichment analysis and motif clustering. Known motifs were annotated against the JASPAR CORE 2026 vertebrate database^121^. Control sequences were generated using the default sequence shuffling strategy in XSTREME. Enriched motifs were ranked according to the statistical significance values reported by XSTREME.

GWAS enrichment analysis was performed for human-specific cCREs in selected cCRE clusters. Single-nucleotide polymorphisms (SNPs) associated with schizophrenia and Alzheimer’s disease were obtained from the NHGRI-EBI GWAS Catalog association table^122^. For each disease *d*, let *S*_*d*_ denote the set of associated SNPs, and let *C*_*q*_ denote cCRE cluster *q*. The observed overlap count was defined as:

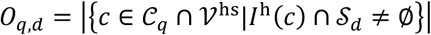

where *C*_hs_ denotes the set of human-specific cCREs and *I*^*h*^(*c*) denotes the human genomic interval of cCRE *c*.

Statistical significance was assessed using 1,000 random permutations against the global human cCRE background. In each permutation, a random set of human cCREs with the same size as *C*_*q*_ ∩ *V*^hs^ was sampled, and its overlap with the corresponding SNPs was computed. Fold enrichment and empirical p-value were calculated by comparing the observed overlap count with the permutation background, as detailed in Supplementary Note 8.

### EpiZoo-Cancer for prioritizing somatic mutations in cancer regulatory contexts

We adapted EpiZoo to pan-cancer scATAC-seq data by incorporating cancer type information into the cell embeddings. The workflow consisted of adaptation to cancer regulatory contexts, LoA score calculation, COSMIC driver-gene enrichment analysis and transcription factor target enrichment analysis.

#### Adaptation to cancer regulatory contexts

We adapted the pretrained EpiZoo model to a pan-cancer scATAC-seq dataset comprising 200,936 cells across eight cancer types^59^. The model retained the human cCRE vocabulary *V*^h^. For each cell from cancer type *a*, we introduced a learnable embedding Emb_cancer_(*a*) to represent the corresponding cancer regulatory context. Given the cell embedding z, extracted from the output vector of the [CLS] token in the MoE transformer, the EpiZoo-Cancer cell embedding was defined as:^93^

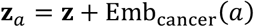

During training, the cancer-type embedding was randomly set to zero with probability 0.1 to encourage the model to retain general regulatory information while learning patterns dependent on cancer contexts. During inference, the full cancer type embedding was used. EpiZoo-Cancer was fine-tuned using the same tasks as EpiZoo pretraining, including cell-cCRE alignment and signal reconstruction.

#### LoA score computation

For each variant *v*, we first identified the human cCRE that contained this variant. The reference and alternative sequence were constructed by replacing the reference allele with the mutated allele. These two sequences were encoded by SEAM to obtain the reference and alternative sequence embeddings, denoted as 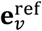 and 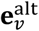, respectively. The sequence perturbation was defined as:

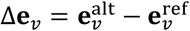

For each cell from the cancer type *a*, the EpiZoo-Cancer cell embedding z_*a*_ was used as the reference embedding, and the perturbed cell embedding was obtained by adding Δe_*v*_. The human-specific signal decoder was then used to predict accessibility profiles before and after the sequence perturbation:

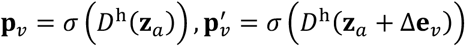

where *D*^h^ denotes the human signal decoder branch, *σ*(·) denotes the sigmoid function. The log fold change vector across the human cCRE vocabulary was computed as:

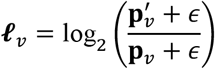

where є is a small pseudocount. The local effect of variant *v* in each cell was defined as the log fold change value at the cCRE that contained this variant. The global impact score was computed for each cell using *L*_2_ norm of the corresponding log fold-change vector across all human cCREs. For variant *v* and cancer type *a*, the 200 cells with the largest global impact scores were selected to construct the sensitive cell set *C*_*a,v*_. The final effect score was defined as the average local effects among these sensitive cells and the LoA score was defined by reversing the sign of the final effect score:

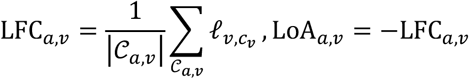

where *c*_*v*_ denotes the human cCRE that contained variant *v*.

#### COSMIC enrichment analysis

Somatic variants from eight cancer types were obtained from TCGA^123^. Variants overlapping human cCREs were retained, yielding 5,959 evaluated noncoding variants. For each cancer type *a*, let *W*_*a*_ denote the evaluated variants from that cancer type. Variants with LoA scores in the top 2.5% were defined as high-impact mutations:

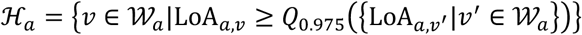

where *Q*_0.975_ denotes the 0.975 quantile.

Each evaluated variant was assigned to its nearest proximal gene using the GENCODE reference annotation^124^. The COSMIC Cancer Gene Census v103 was used as the reference cancer driver gene catalog, and only Tier 1 genes were retained. For each cancer type, Fisher’s exact test was used to assess whether high-impact mutations were assigned to COSMIC Tier 1 driver genes compared with the remaining evaluated variants. The contingency table and odds ratio calculation are provided in Supplementary Note 9.

#### Transcription factor target enrichment analysis

Transcription factor target enrichment analysis was performed using Metascape. High-impact mutations prioritized by the zero-shot and fine-tuned EpiZoo-Cancer were separately assigned to their nearest proximal genes using the GENCODE reference annotation. The resulting gene lists were submitted to Metascape using the multiple gene list mode. Enrichment analysis was performed using the “Transcription Factor Targets” category with the default background. Enriched transcription factor target terms were ranked according to the enrichment statistics reported by Metascape and visualized for comparison between the zero-shot and fine-tuned models.

### Cell-type-specific chromatin accessibility prediction and downstream analysis

EpiZoo was used to predict cell-type-specific chromatin accessibility from DNA sequence and learned cell embeddings. The workflow included model training, nucleotide-resolution in silico attribution, and motif annotation of sequence segments with high attribution scores.

#### Model training

We adopted a chromosomal based split following the CREsted benchmark. In the BICCN2 dataset, chromosomes 8 and 10 were reserved for validation, chromosomes 9 and 18 were reserved for testing, and the remaining chromosomes were used for training.

For each cell type *τ*, a cell-type embedding z_*τ*_ was obtained by averaging EpiZoo cell embeddings across cells assigned to that cell type. To predict the accessibility of a peak *p* under the context of cell type *τ*, the SEAM-derived sequence embedding of *p* was concatenated with the cell-type embedding:

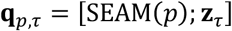

where [·;·] denotes vector concatenation. The concatenated representation was passed to a two-layer fusion MLP with a Softplus output. The model was optimized using a Cosine-MSE Log Loss, with detailed definition provided in Supplementary Note 10.

The model training was divided into three stages. In the warm-up stage, the model was trained on consensus peaks for one epoch with SEAM frozen, and only the fusion MLP was updated. In the joint optimization stage, SEAM was unfrozen and updated together with the fusion MLP on consensus peaks. In the specialization stage, the model was fine-tuned on cell-type-specific peaks. Model performance was evaluated on test peaks using PCCs, as detailed in Supplementary Note 10. **Nucleotide-resolution in silico attribution**. Nucleotide-level contribution scores were obtained using integrated gradients^125^ computed with respect to the continuous DNABERT2 input embeddings. For each target cell type, the corresponding cell-type embedding was fixed during attribution, and gradients were computed only with respect to sequence input embeddings. Token-level attribution scores were projected back to nucleotide positions for visualization as attribution logos. Detailed attribution formulas and mapping procedures from token to nucleotide are provided in Supplementary Note 10.

#### Motif annotation

We extracted sequence segments with high attribution scores from the AiE2428m and AiE2587m enhancer regions. These segments were used as candidate query motifs and compared against known transcription factor motifs using Tomtom^126^ from the MEME Suite online server. Motif similarity search was performed against the JASPAR CORE 2026 vertebrate motif database. The matched transcription factor motifs reported by Tomtom were used as candidate annotations.

## Data availability

The pretraining data were collected from 42 published datasets^74-113^ and publicly available datasets from the 10x Genomics website (https://www.10xgenomics.com/datasets/). Four additional published datasets^13,28-30^ were used for benchmarking. Detailed information on all pretraining and benchmark datasets, including their sources, accession numbers or download links, species, tissues, sequencing technologies and summary statistics, is provided in Supplementary Table 1. The datasets used to develop and evaluate EpiZoo across four additional species were also obtained from published studies^8,45,46^, with detailed information provided in Supplementary Table 3. The data used for EpiZoo-Cancer are available from the TCGA Publication Page at https://gdc.cancer.gov/about-data/publications/TCGA-ATAC-Seq-2024. The mouse BICCN2 data were obtained from the processed datasets provided by CREsted^70^ and are available at https://resources.aertslab.org/CREsted/manuscript_data.

## Code availability

The source code and analysis scripts for EpiZoo are freely available on GitHub at https://github.com/likeyi19/EpiZoo.

## Acknowledgements

This work was supported by the National Key Research and Development Program of China, grant numbers 2025YFC3409300 (R.J.), 2023YFF1204802 (R.J.) the National Natural Science Foundation of China, grant numbers 32550616 (R.J.), 62273194 (R.J.) and Beijing Natural Science Foundation, grant number L242026 (R.J.).

## Author information

### Contributions

R.J. and X.C. conceived the study and supervised the project. K.L. and X.C. collected and processed all data used in Omni-scATAC and the downstream analyses. X.C. designed the EpiZoo framework, and K.L. performed the pretraining of EpiZoo. K.L. and X.C. designed and implemented the downstream analyses and validation experiments. Q.J. contributed to the analysis of unsupervised feature extraction results. Z.W. contributed to the analysis of cell type annotation results. K.L., X.C. and R.J. wrote and revised the manuscript, with input from all authors.

