## Supplementary Notes for "EpiZoo: a DNA sequence-aware foundation model for cross-species single-cell epigenomics"

Contents

**Supplementary Notes** 4

**Supplementary Note 1: Ablation analysis of DNA sequence and cCRE identity embeddings** 4

**Supplementary Note 2: GREAT analysis on the EpiZoo-imputed data and raw data** 5

**Supplementary Note 3: cCRE construction, filtering and assessment** 6

**Supplementary Note 4: Global TF-IDF calculation** 7

**Supplementary Note 5: LoRA formulations and ablation settings** 8

**Supplementary Note 6: Evaluation metrics for single-cell analysis** 9

**Supplementary Note 7: Analysis of EpiZoo-Evo cell embeddings** 11

**Supplementary Note 8: Analysis of EpiZoo-Evo cCRE embeddings** 13

**Supplementary Note 9: COSMIC enrichment analysis of EpiZoo-Cancer** 14

**Supplementary Note 10: Cell-type-specific chromatin accessibility prediction and analysis** 15

**Supplementary Figures** 17

**Supplementary Figure 1** 17

**Supplementary Figure 2** 18

**Supplementary Figure 3** 19

**Supplementary Figure 4** 20

**Supplementary Figure 5** 21

**Supplementary Figure 6** 22

**Supplementary Figure 7** 23

**Supplementary Figure 8** 24

**Supplementary Figure 9** 25

**Supplementary Figure 10** 26

**Supplementary Figure 11** 27

**Supplementary Figure 12** 28

**Supplementary Figure 13** 29

**Supplementary Figure 14** 30

**Supplementary Figure 15** 31

**Supplementary Figure 16** 32

**Supplementary Figure 17** .......... 33

### Supplementary Notes

#### Supplementary Note 1: Ablation analysis of DNA sequence and cCRE identity embeddings

To assess the contributions of the DNA sequence embedding and cCRE identity embedding in EpiZoo, we performed an ablation analysis on the four benchmark datasets. We compared three model configurations under the same fine-tuning and evaluation framework. The full EpiZoo model used both SEAM-derived DNA sequence embeddings and learnable cCRE identity embeddings. The DNA sequence-only variant retained only the sequence embeddings, whereas the cCRE identity-only variant retained only the identity embeddings. In all three configurations, rank embeddings were retained to capture importance based on accessibility. Apart from the ablated embedding modules, the remaining trainable modules, including the MoE transformer and species-specific signal decoders, were fine-tune using the same procedure.

The DNA sequence-only variant evaluated the contribution of regulatory priors, because the sequence embeddings were precomputed and remained frozen during downstream fine-tuning. The cCRE identity-only variant evaluated the contribution of the epigenomic context in absence of the DNA sequence information. The full EpiZoo model consistently achieves the strongest clustering performance across the benchmark datasets, whereas the cCRE identity-only variant outperforms the DNA sequence-only variant but remains below the full model. These results indicate that the integration of DNA sequence information and sequence-independent epigenomic context allows EpiZoo to learn cell representations that are more informative than those produced by either component alone.

**Supplementary Note 2: GREAT analysis on the EpiZoo-imputed data and raw data**

We performed GREAT analysis[1] using cell-type-specific cCREs from the Kanemaru2023 dataset. Raw data refers to the dropout matrix after global TF-IDF transformation and before EpiZoo imputation. Both the raw data and the imputed data were restricted to the same filtered cCRE set used in the data imputation task. For each matrix, we ranked cCREs using the 'sc.tl.rank_genes_groups' function and the top 5,000 ranked cCREs for each cell type were submitted to GREAT for enrichment analysis. The top five enriched biological processes were ranked by the binomial FDR q-value. Relevant Biological processes were highlighted based on their functional relationship to the corresponding cell type (Supplementary Figure 15).

For atrial cardiomyocytes, both raw and imputed data recovered biological processes related to muscle cells[2]. Notably, EpiZoo imputation substantially increased the enrichment significance of these functionally relevant biological processes. For example, myofibril assembly increased from $5.40 \times 10^{-64}$ in the raw data to $1.56 \times 10^{-111}$ after EpiZoo imputation, and sarcomere organization increased from $1.74 \times 10^{-59}$ to $4.18 \times 10^{-106}$. These results indicate that EpiZoo strengthens the recovery of cardiomyocyte regulatory programs rather than merely preserving the baseline signal.

For endothelial cells, EpiZoo imputation improved both the biological specificity and enrichment strength of the recovered biological processes. In the raw data, the top enriched biological processes included endothelium development and endothelial cell differentiation, but also less specific terms related to type I interferon response and host interaction. By contrast, the EpiZoo-imputed data recovered angiogenesis, a core process of new blood vessel formation[3], as the top enriched biological process, together with endothelium development and endothelial cell differentiation. These results demonstrate that the imputed data generated by EpiZoo provide a more functionally informative representation of endothelial regulatory programs.

Together, this analysis supports that EpiZoo imputation improves not only clustering performance and quantitative signal recovery, but also the biological interpretability of downstream enrichment analysis.

### Supplementary Note 3: cCRE construction, filtering and assessment

The scATAC-seq datasets were represented within species-specific cCRE vocabularies. For human, we adopted the cCRE vocabulary established in the Human-scATAC-Corpus[4]. For mouse, we used the cCRE vocabulary curated by the Lu2025 dataset[5], a large-scale mouse scATAC-seq atlas spanning diverse tissues and biological contexts. Public datasets provided as fragment files or peak-by-cell matrices were harmonized into cell-by-cCRE matrices over cCRE vocabulary of the corresponding species. The coverage of the human cCRE vocabulary has been systematically evaluated in EpiAgent[6] and Human-scATAC-Corpus[4]. We therefore applied the same overlap assessment to the mouse cCRE vocabulary.

We compared the mouse cCRE vocabulary with independently generated peak or cCRE collections from other mouse datasets. Genomic overlaps were calculated between the mouse cCRE vocabulary and each external collection, and the overlap proportions were summarized across datasets. The substantial overlap observed across independent datasets supported the broad coverage of the mouse cCRE vocabulary (Supplementary Figure 17).

We further filtered cCREs according to their accessibility across the whole corpus. For each cCRE, we counted the total number of cells in which it was accessible across datasets of the corresponding species. We then inspected the empirical complementary cumulative distribution of these counts. Based on the empirical curves, we retained human cCREs accessible in at least 15,000 cells and mouse cCREs accessible in at least 20,000 cells. The resulting filtered cCREs were used for tokenization, pretraining, fine-tuning and downstream analysis.

**Supplementary Note 4: Global TF-IDF calculation**

For each dataset $d$ of species $s$, let

$$\mathbf{X}_d \in \mathbb{R}^{N_d \times |\mathcal{V}^s|}$$

denote its raw cell-by-cCRE matrix, where $x_{d,ij}$ denotes the count value of cCRE $j$ in cell $i$, $N_d$ denotes the cell number and $\mathcal{V}^s$ denotes the cCRE vocabulary. Let $\mathcal{D}$ denote the set of datasets of the corresponding species. The document frequency of cCRE $j$ was calculated as:

$$df_j = \sum_{d \in \mathcal{D}} \frac{1}{N_d} \sum_{i=1}^{N_d} x_{d,ij}$$

The document frequency vector $\mathbf{df}^s = \left(df_1, df_2, \ldots, df_{|\mathcal{V}^s|}\right)^{\mathrm{T}}$ was computed once from the corresponding species-level reference corpus and then kept fixed. The row-normalized accessibility of cCRE $j$ in cell $i$ was calculated as:

$$tf_{d,ij}^s = \frac{x_{d,ij}}{\sum_{c \in \mathcal{V}^s} x_{d,ic}}$$

Inverse document-frequency normalization was then applied using the fixed $\mathbf{df}^s$:

$$u_{d,ij}^s = \frac{tf_{d,ij}^s}{df_j}$$

The final TF-IDF score of cCRE $j$ in cell $i$ was calculated as:

$$tfidf_{d,ij} = \log\left(1 + \gamma u_{d,ij}^s\right)$$

where $\gamma = 10{,}000$ is the scaling factor.

**Supplementary Note 5: LoRA formulations and ablation settings**

**LoRA for embedding matrices**

For a learnable embedding matrix $\mathbf{E} \in \mathbb{R}^{V\times d}$, where $V$ denotes the vocabulary size and $d$ denotes the hidden size, LoRA introduced a low-rank update as:

$$\Delta\mathbf{E} = \mathbf{AB}$$

where $\mathbf{A} \in \mathbb{R}^{V\times r}$, $\mathbf{B} \in \mathbb{R}^{r\times d}$, and $r \ll d$. The adapted embedding was computed as:

$$\mathbf{E}' = \mathbf{E} + \frac{\alpha}{r}\Delta\mathbf{E}$$

where $\alpha$ is a scaling factor. The pretrained embedding $\mathbf{E}$ was kept frozen, whereas $\mathbf{A}$ and $\mathbf{B}$ were trainable. Matrix $\mathbf{A}$ was initialized by Kaiming uniform initialization and $\mathbf{B}$ was initialized to zero.

**LoRA for linear layers**

For a pretrained linear layer weight matrix $\mathbf{W} \in \mathbb{R}^{d_{in}\times d_{out}}$, LoRA introduced a low-rank update as:

$$\mathbf{W}' = \mathbf{W} + \frac{\alpha}{r}\mathbf{AB}$$

where $\mathbf{A} \in \mathbb{R}^{d_{in}\times r}$ and $\mathbf{B} \in \mathbb{R}^{r\times d_{out}}$ denote the trainable low-rank matrices. The pretrained weight $\mathbf{W}$ was frozen during LoRA. LoRA was applied to query, key, value and output projection matrices in the multi-head self-attention blocks, linear layers within the MoE networks, and linear layers in the signal decoder.

**Ablation of fine-tuning strategies**

We evaluated eight configurations by assigning either FFT or LoRA to each of the three core modules: the DNA sequence-aware embedding module, the MoE transformer and the species-specific signal decoders.

1. FFT baseline: all pretrained weights were unfrozen and updated.
2. Single-module LoRA: LoRA was applied to one of the three modules, whereas the other two modules underwent FFT.
3. Dual-module LoRA: LoRA was applied to two modules, whereas the remaining module underwent FFT.
4. Full LoRA: LoRA was applied to all three modules, with original pretrained weights frozen.

### Supplementary Note 6: Evaluation metrics for single-cell analysis

#### Clustering metrics

Louvain clustering was performed on the cell embeddings, and the resulting cluster assignments were compared with the cell labels using normalized mutual information (NMI) and adjusted rand index (ARI).

Let $U = \{u_1, \dots, u_R\}$ denote the set of predicted clusters and $V = \{v_1, \dots, v_C\}$ denote the set of reference cell-type labels. Let $n_{rc}$ be the number of cells shared by predicted cluster $u_r$ and reference label $v_c$, $n_{r\cdot} = \sum_c n_{rc}$ and $n_{\cdot c} = \sum_r n_{rc}$ be the corresponding marginal counts, and $n$ be the total number of cells. The mutual information between $U$ and $V$ was calculated as:

$$I(U;V) = \sum_{r=1}^{R}\sum_{c=1}^{C}\frac{n_{rc}}{n}\log\frac{n \cdot n_{rc}}{n_{r\cdot} \cdot n_{\cdot c}}$$

where terms with $n_{rc} = 0$ were omitted.

The entropies of U and V were calculated as:

$$H(U) = -\sum_{r=1}^{R}\frac{n_{r\cdot}}{n}\log\frac{n_{r\cdot}}{n}, H(V) = -\sum_{c=1}^{C}\frac{n_{\cdot c}}{n}\log\frac{n_{\cdot c}}{n}$$

NMI was defined as:

$$\mathrm{NMI}(U,V) = \frac{2I(U;V)}{H(U) + H(V)}$$

ARI was defined as:

$$\mathrm{ARI} = \frac{\sum_{ij}\binom{n_{ij}}{2} - \frac{\sum_i\binom{a_i}{2}\sum_j\binom{b_j}{2}}{\binom{n}{2}}}{\frac{1}{2}\left[\sum_i\binom{a_i}{2} + \sum_j\binom{b_j}{2}\right] - \frac{\sum_i\binom{a_i}{2}\sum_j\binom{b_j}{2}}{\binom{n}{2}}}$$

where $n_{ij}$ denotes the number of cells shared by predicted cluster $i$ and reference label $j$, $a_i$ and $b_j$ are the marginal sums, and $n$ is the total number of cells. Higher NMI and ARI values indicate stronger agreement between predicted clusters and reference cell-type labels.

#### Classification metrics

Classification performance was evaluated using accuracy, Cohen's kappa coefficient (κ) and macro F1 (mF1) score. Accuracy was defined as:

$$\mathrm{acc} = \frac{1}{n}\sum_i \mathbb{I}(y_i = \hat{y}_i)$$

where $y_i$ and $\hat{y}_i$ denote the true and predicted labels of cell $i$ and $\mathbb{I}(\cdot)$ denotes the indicator function.

κ was computed as:

$$\kappa = \frac{p_o - p_e}{1 - p_e}$$

where $p_o$ is the observed agreement and $p_e$ is the expected agreement under random label assignment estimated from the marginal label distributions. Specifically,

$$p_o = \frac{1}{n}\sum_i \mathbb{I}(y_i = \hat{y}_i)\,, p_e = \sum_{k=1}^{K}\left(\frac{n_{k\cdot}}{n}\right)\left(\frac{n_{\cdot k}}{n}\right)$$

where $n_{k\cdot}$ denotes the number of cells whose true label is cell type $k$, and $n_{\cdot k}$ denotes the number of cells predicted as cell type $k$.

For each cell type $k$, precision and recall were computed as:

$$\text{Precision}_k = \frac{\text{TP}_k}{\text{TP}_k + \text{FP}_k}, \text{Recall}_k = \frac{\text{TP}_k}{\text{TP}_k + \text{FN}_k}$$

where $\text{TP}_k$, $\text{FP}_k$ and $\text{FN}_k$ denote the numbers of true-positive, false-positive and false-negative predictions for cell type $k$, respectively. The F1 score for cell type $k$ was defined as:

$$\text{F1}_k = \frac{2 \cdot \text{Precision}_k \cdot \text{Recall}_k}{\text{Precision}_k + \text{Recall}_k}$$

The mF1 score was then calculated as the unweighted average of class-wise F1 scores across all cell types:

$$\text{mF1} = \frac{1}{K}\sum_{k=1}^{K} \text{F1}_k$$

**Pearson correlation coefficient**

For two vectors $\mathbf{x}$ and $\mathbf{y}$, Pearson correlation coefficient (PCC) was calculated as:

$$\text{pearson}(\mathbf{x}, \mathbf{y}) = \frac{\sum_i (x_i - \bar{\mathbf{x}})(y_i - \bar{\mathbf{y}})}{\sqrt{\sum_i (x_i - \bar{\mathbf{x}})^2}\sqrt{\sum_i (y_i - \bar{\mathbf{y}})^2}}$$

**Supplementary Note 7: Analysis of EpiZoo-Evo cell embeddings**

**Harmony integration**

Harmony integration[7] was applied to EpiZoo-Evo cell embeddings to reduce batch effects associated with species during joint visualization of human and macaque cells. Harmony was performed on the raw EpiZoo-Evo cell embedding matrix using species identity as the batch covariate.

Let

$$\mathbf{H} = [\mathbf{h}_1, \mathbf{h}_2, \dots, \mathbf{h}_N]^{\mathrm{T}} \in \mathbb{R}^{n \times d}$$

denote the raw EpiZoo-Evo cell embedding matrix, where $\mathbf{h}_i$ denote the cell embedding of cell $i$, $N$ is the number of cells and $d$ is the embedding dimension. Let $b_i \in \{\mathrm{h}, \mathrm{m}\}$ denote the species label of cell $i$, corresponding to human or macaque. Harmony estimates species-associated correction terms in the embedding space and produces a corrected embedding matrix

$$\mathbf{H}^{\mathrm{harm}} = \left[\mathbf{h}_1^{\mathrm{harm}}, \mathbf{h}_2^{\mathrm{harm}}, \dots, \mathbf{h}_N^{\mathrm{harm}}\right]^{\mathrm{T}}$$

Following the Harmony framework, each corrected cell embedding can be written as

$$\mathbf{h}_i^{\mathrm{harm}} = \mathbf{h}_i - \sum_{k=1}^{K} r_{ik} \boldsymbol{\delta}_{k,b_i}$$

where $r_{ik}$ denotes the soft assignment of cell $i$ to Harmony cluster $k$, and $\boldsymbol{\delta}_{k,b_i} \in \mathbb{R}^d$ denotes the species-specific correction vector estimated for cluster $k$ and species $b_i$. Harmony iteratively estimates the soft cluster assignments and correction vectors until convergence. The corrected embeddings $\mathbf{H}^{\mathrm{harm}}$ were used for joint UMAP visualization of human and macaque cells. Cross-species label transfer was performed using the raw, unintegrated EpiZoo-Evo cell embeddings rather than the embeddings corrected by Harmony.

**Cross-species label transfer**

Let $\mathbf{E}^s \in \mathbb{R}^{N^s \times d}$ denote the raw embedding matrix for species $s$, where $N^s$ denotes the number of cells and $d$ denotes the embedding dimension. For each species, the embedding of cell $i$ were first mean-centered:

$$\tilde{\mathbf{e}}_i^s = \mathbf{e}_i^s - \frac{1}{N^s} \sum_{j=1}^{N^s} \mathbf{e}_j^s$$

The centered embeddings were then L2-normalized:

$$\bar{\mathbf{e}}_i^s = \frac{\tilde{\mathbf{e}}_i^s}{\left\|\tilde{\mathbf{e}}_i^s\right\|_2}$$

For a query cell $j$ from the target species $s'$ and a reference cell $i$ from species $s$, cosine similarity was computed as:

$$\mathrm{sim}(i,j) = \bar{\mathbf{e}}_j^{s'} \cdot \bar{\mathbf{e}}_i^s$$

The top $k$ nearest reference cells were selected according to cosine similarity ($k = 20$). The predicted label of query cell $j$ was assigned as:

$$\hat{y}_j = \mathrm{mode}\left(\{y_q | q \in \mathrm{neighbor}_k(j)\}\right)$$

where $\mathrm{mode}(\cdot)$ denotes the majority voting function, $\mathrm{neighbor}_k(\cdot)$ denotes the set of top $k$ nearest reference cells ranked by cosine similarity.

The overall label transfer accuracy was calculated as:

$$\text{acc} = \frac{1}{N}\sum_{j=1}^{N} \mathbb{I}(y_j = \hat{y}_j)$$

where $N$ denotes the number of query cells. For each cell type, accuracy was calculated as the fraction of correctly predicted cells among all query cells belonging to that cell type.

### Supplementary Note 8: Analysis of EpiZoo-Evo cCRE embeddings

#### Homologous gene mapping enrichment

For C4 macro-cCRE analysis, embedding-based human-macaque cCRE pairs were compared with randomly paired human-specific and macaque-specific cCREs. Each cCRE was assigned to its nearest proximal gene, and a pair was considered homologous if the assigned human and macaque genes were annotated as homologous genes.

The contingency table was defined as:

| | Homologous | Not homologous |
|---|---|---|
| EpiZoo pairs | $a$ | $b$ |
| Random pairs | $c$ | $d$ |

The odds ratio was calculated as:

$$\mathrm{OR} = \frac{ad}{bc}$$

A one-sided Fisher's exact test was used to evaluate whether cCRE pairs based on embeddings showed a higher frequency of homologous gene matches than randomly paired cCREs.

#### GWAS enrichment analysis

For each cluster and disease pair, we conducted $B = 1000$ random permutations. Let $O_{q,d}$ denote the observed number of overlaps between human-specific cCREs in the evaluated cluster and disease-associated SNPs. In permutation $b$, the random overlap count was denoted as $R_{q,d}^{(b)}$. The expected overlap count was calculated as:

$$E_{q,d} = \frac{1}{B}\sum_{b=1}^{B} R_{q,d}^{(b)}$$

Fold enrichment and empirical p-value were calculated as:

$$FE_{q,d} = \frac{O_{q,d}}{E_{q,d}}$$

$$p_{q,d} = \frac{1 + \sum_{b=1}^{B} \mathbb{I}\left(R_{q,d}^{(b)} \geq O_{q,d}\right)}{B+1}$$

This procedure was repeated for each selected cCRE cluster and disease.

**Supplementary Note 9: COSMIC enrichment analysis of EpiZoo-Cancer**

For each evaluated mutation, the nearest proximal gene was identified using GENCODE annotation. Each variant was then annotated according to whether its assigned gene belonged to the COSMIC Cancer Gene Census Tier 1 driver list.

For each cancer type, the contingency table was defined as:

| | Mapped | Not mapped |
|---|---|---|
| High-impact mutation | $a$ | $b$ |
| Not high-impact mutation | $c$ | $d$ |

The odds ratio was calculated as:

$$\mathrm{OR} = \frac{ad}{bc}$$

Fisher's exact test was used to evaluate whether high-impact mutations were preferentially assigned to COSMIC Tier 1 driver genes.

**Supplementary Note 10: Cell-type-specific chromatin accessibility prediction and analysis**

**Cosine-MSE Log Loss**

Let $\hat{y}_{p,\tau}$ and $y_{p,\tau}$ denote the predicted and observed accessibility values of peak $p$ in cell type $\tau$, respectively. The log-space mean squared error was defined as:

$$\mathcal{L}_{\text{MSE}} = \frac{1}{C}\sum_{\tau=1}^{C}\left(\log\left(1+\mu\hat{y}_{p,\tau}\right) - \log\left(1+\mu y_{p,\tau}\right)\right)^2$$

where $\mu = 1000$ and $C$ denotes the number of cell types. The cosine similarity between predicted and observed accessibility profiles was computed as:

$$\text{Cos}\left(\hat{\mathbf{y}}_p, \mathbf{y}_p\right) = \frac{\hat{\mathbf{y}}_p \cdot \mathbf{y}_p}{\| \hat{\mathbf{y}}_p \|_2 \| \mathbf{y}_p \|_2 + \epsilon}$$

where $\epsilon$ is a small pseudocount. The final training objective was:

$$\mathcal{L}_{\text{seq}} = \mathcal{L}_{\text{MSE}} + w\left(1 - \text{Cos}\left(\hat{\mathbf{y}}_p, \mathbf{y}_\mathbf{p}\right)\right)$$

where $w$ denotes the weighting hyperparameter.

**Performance evaluation**

Before correlation calculation, predicted and observed values were clamped at zero and transformed as:

$$T(x) = \log(1 + \mu \max(x, 0))$$

where $\mu$ denotes the scaling multiplier. Cell-type-level PCC was computed across test peaks for each cell type:

$$r_\tau = \text{pearson}\left(\left\{T\left(\hat{y}_{p,\tau}\right)\right\}_{p\in\mathcal{P}_{\text{test}}}, \left\{T\left(y_{p,\tau}\right)\right\}_{p\in\mathcal{P}_{\text{test}}}\right)$$

where $\mathcal{P}_{\text{test}}$ denotes the held-out test peak set. Peak-level PCC was computed across cell types for each test peak:

$$r_p = \text{pearson}\left(\left\{T\left(\hat{y}_{p,\tau}\right)\right\}_{\tau=1}^{C}, \left\{T\left(y_{p,\tau}\right)\right\}_{\tau=1}^{C}\right)$$

where $C$ denotes the number of cell types. The Pearson correlation function $\text{pearson}(\cdot,\cdot)$ is defined in Supplementary Note 6.

**Integrated gradients attribution**

Nucleotide-level contribution scores were computed using integrated gradients with respect to the continuous DNABERT2 input embeddings. Let

$$\mathbf{E} \in \mathbb{R}^{L\times d_{\text{tok}}}$$

denote the DNABERT2 token embedding matrix, where $L$ is the tokenized sequence length and $d_{\text{tok}} = 768$. The target cell-type embedding was fixed during attribution. Let $F_\tau(\mathbf{E})$ denote the scalar accessibility score predicted for cell type $\tau$, and let $\mathbf{Z}$ denote a zero embedding baseline. For token $t$ and embedding dimension $r$, the attribution was computed as:

$$A_{t,r} = \left(E_{t,r} - Z_{t,r}\right) \times \frac{1}{M}\sum_{m=1}^{M}\frac{\partial F_\tau\left(\mathbf{Z} + \frac{m}{M}(\mathbf{E} - \mathbf{Z})\right)}{\partial E_{t,r}}$$

where $M = 50$ denotes the number of interpolation steps. The token-level attribution score was obtained by summing across the embedding dimension:

$$a_t = \sum_{r=1}^{d_{\text{tok}}} A_{t,r}$$

Special and uninformative tokens were excluded when attribution scores were mapped to nucleotides. For each remaining subword token produced by the tokenizer, tokenizer prefixes were removed, and the attribution score of that token was assigned to the corresponding nucleotides in the original sequence. The resulting attribution profile at the base level was smoothed using a one-dimensional Gaussian filter with $\sigma = 1$ and visualized as an attribution logo at nucleotide resolution.

### Supplementary Figures

#### Supplementary Figure 1

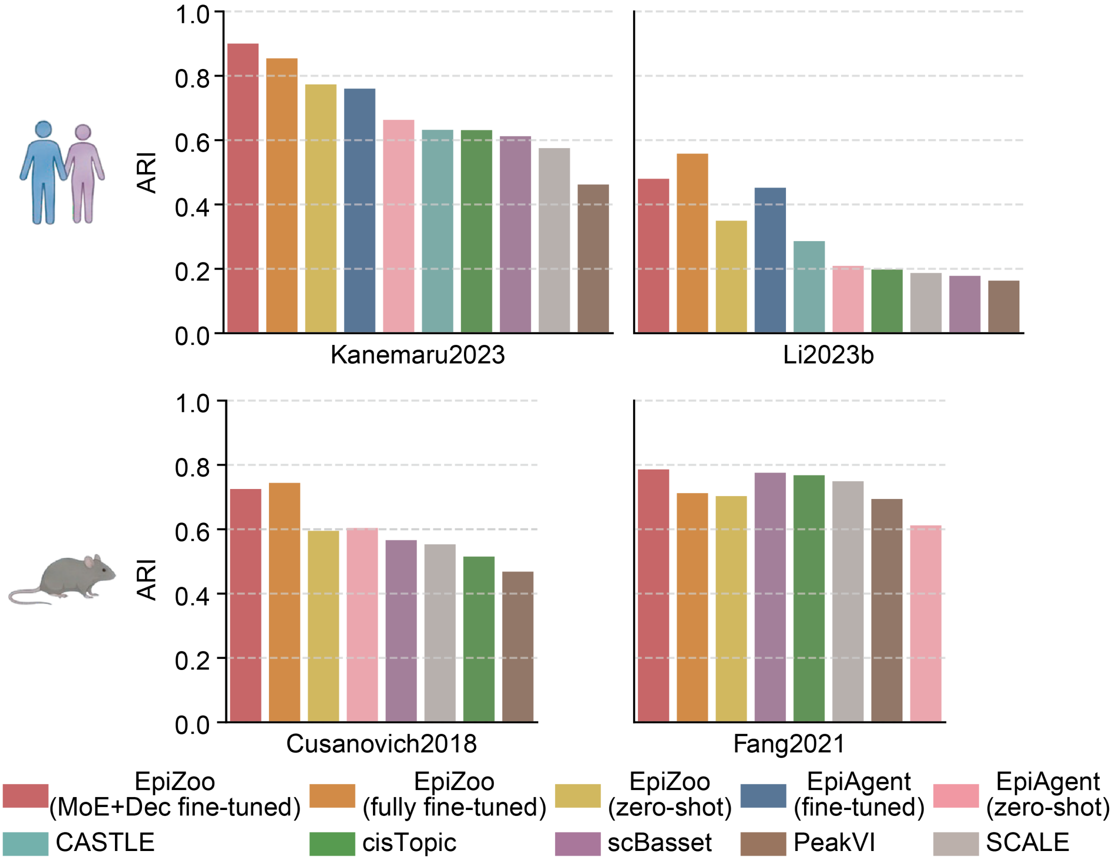


**Supplementary Figure 1.** Quantitative evaluation of unsupervised Louvain clustering performance across four datasets.

**Supplementary Figure 2**

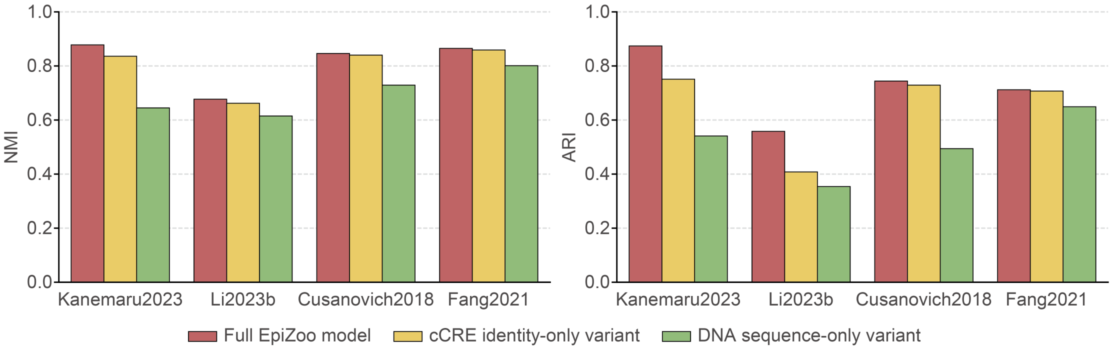


**Supplementary Figure 2.** Clustering performance in the embedding ablation analysis across four datasets.

### Supplementary Figure 3

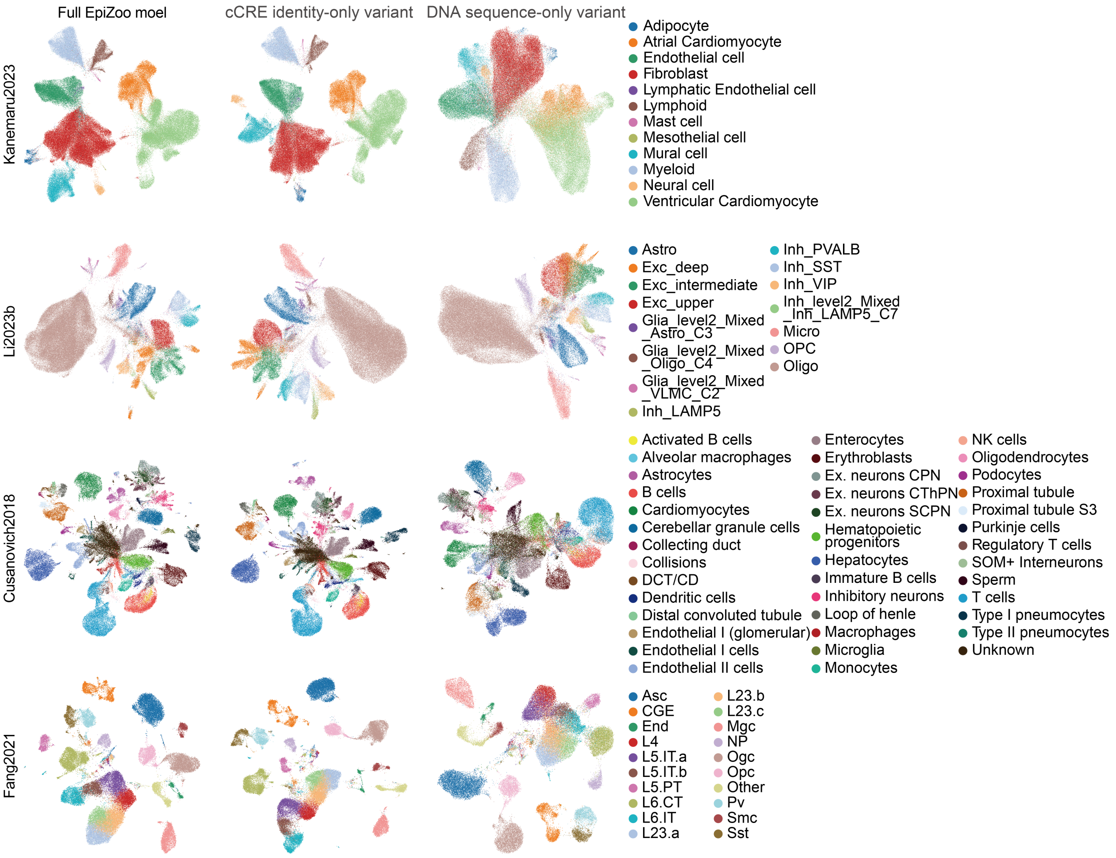


**Supplementary Figure 3.** UMAP visualizations in the embedding ablation analysis across four datasets.

### Supplementary Figure 4

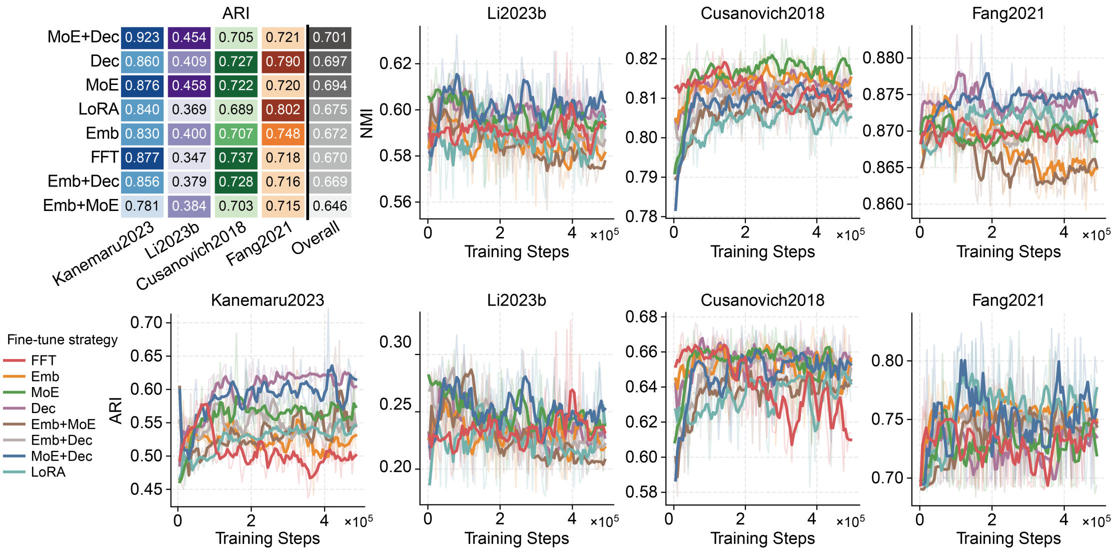


**Supplementary Figure 4.** Systematic evaluation of parameter-efficient fine-tuning strategies.

### Supplementary Figure 5

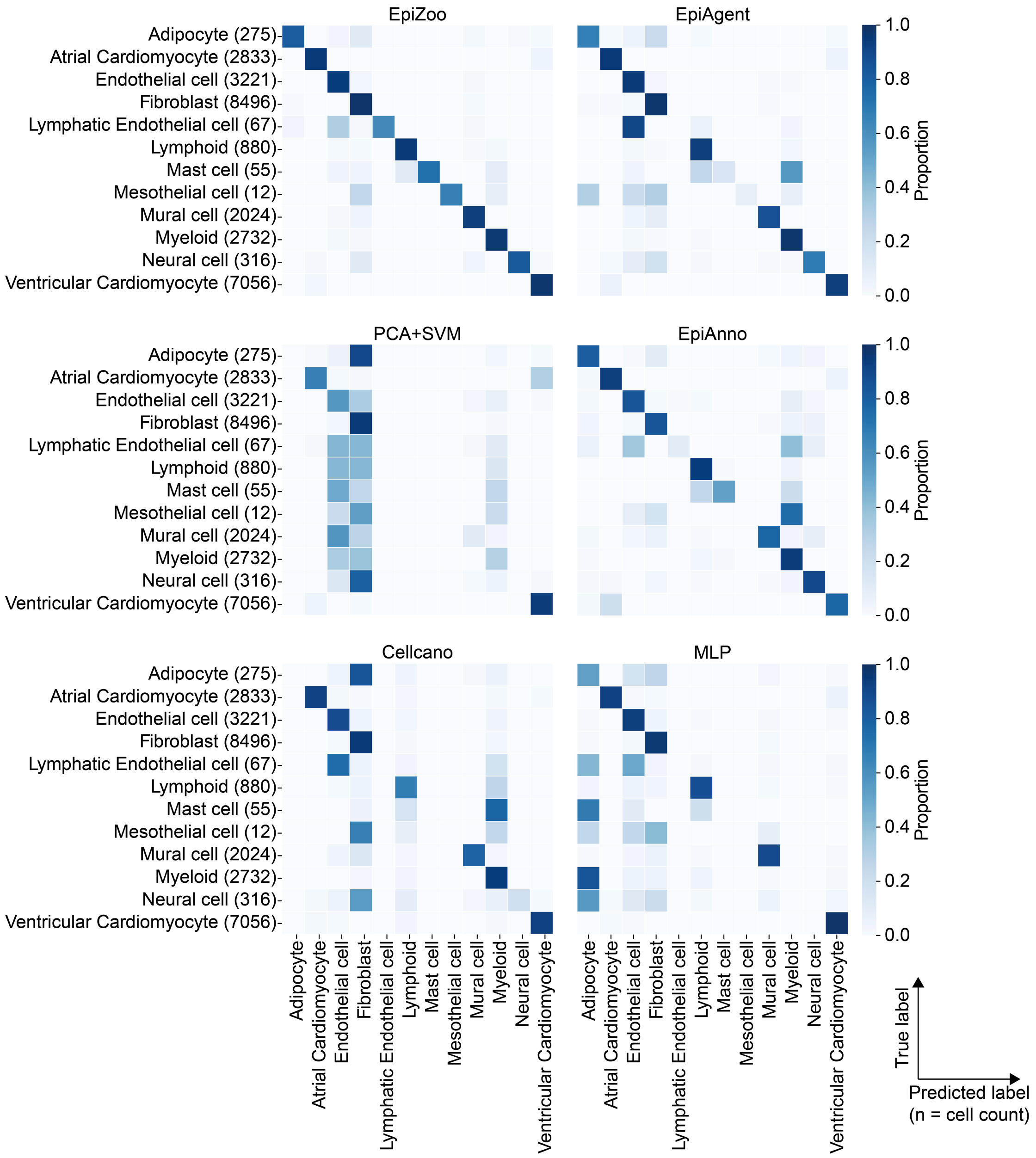


**Supplementary Figure 5.** Normalized confusion matrices of the Kanemaru2023 dataset in cell-type annotation based on the intra-dataset five-fold cross-validation. Rows represent true cell types, and columns represent predicted cell types. Numbers in parentheses indicate the number of cells in each cell type.

### Supplementary Figure 6

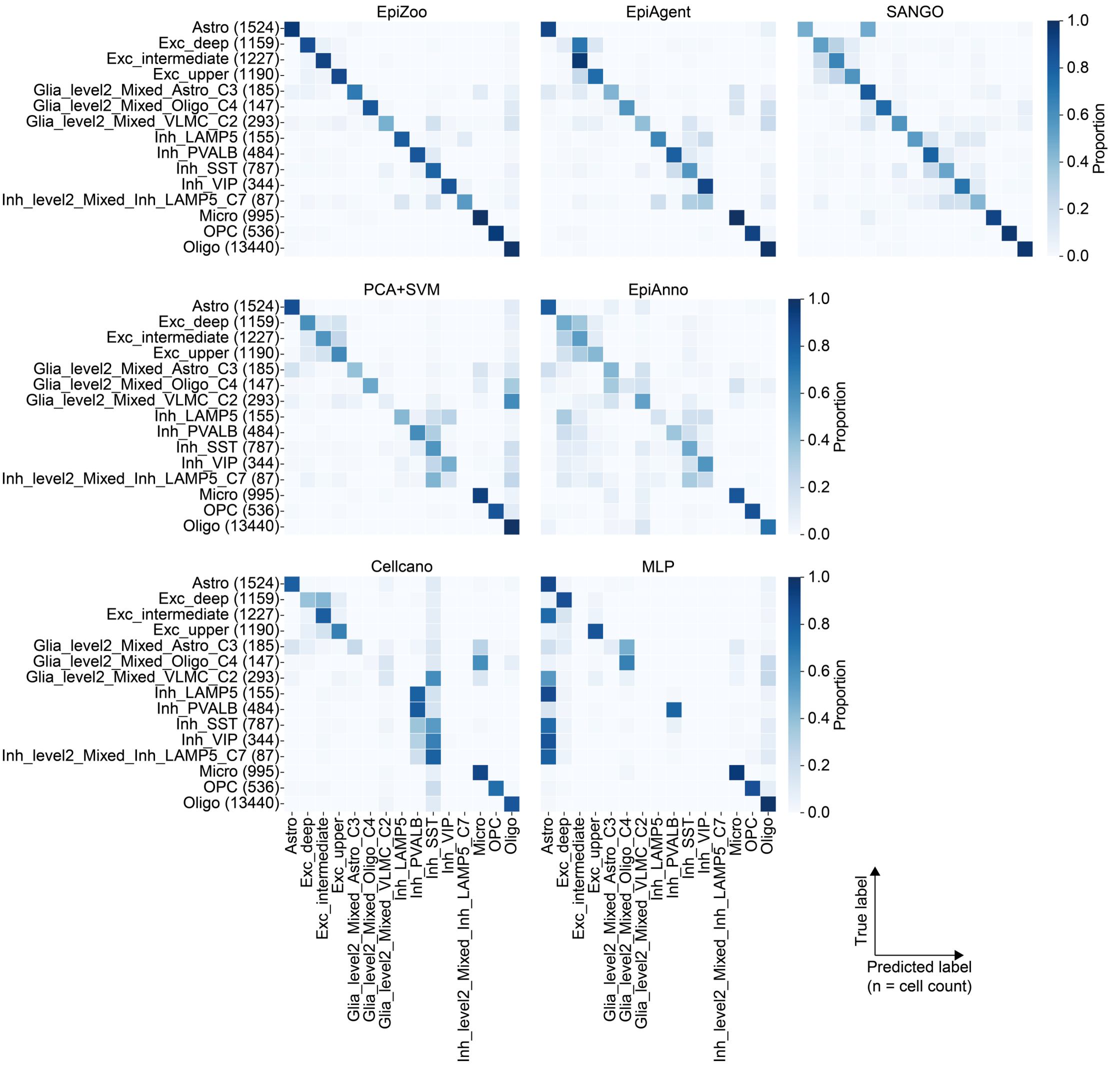


**Supplementary Figure 6.** Normalized confusion matrices of the Li2023b dataset in cell-type annotation based on the intra-dataset five-fold cross-validation. Rows represent true cell types, and columns represent predicted cell types. Numbers in parentheses indicate the number of cells in each cell type.

**Supplementary Figure 7**

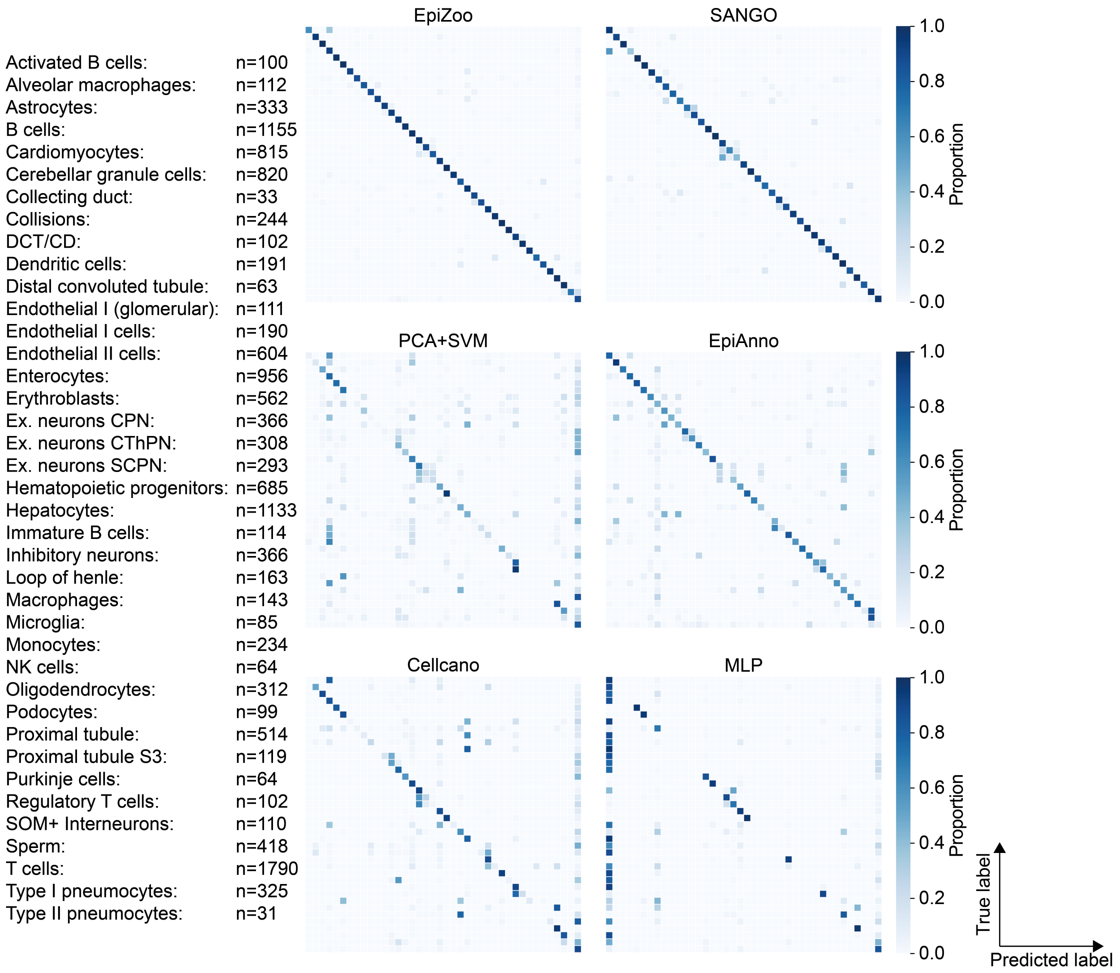


**Supplementary Figure 7.** Normalized confusion matrices of the Cusanovich2018 dataset in cell-type annotation based on the intra-dataset five-fold cross-validation. Rows represent true cell types, and columns represent predicted cell types. Cell-type labels and corresponding cell counts shown on the left, aligning with the matrix rows.

**Supplementary Figure 8**

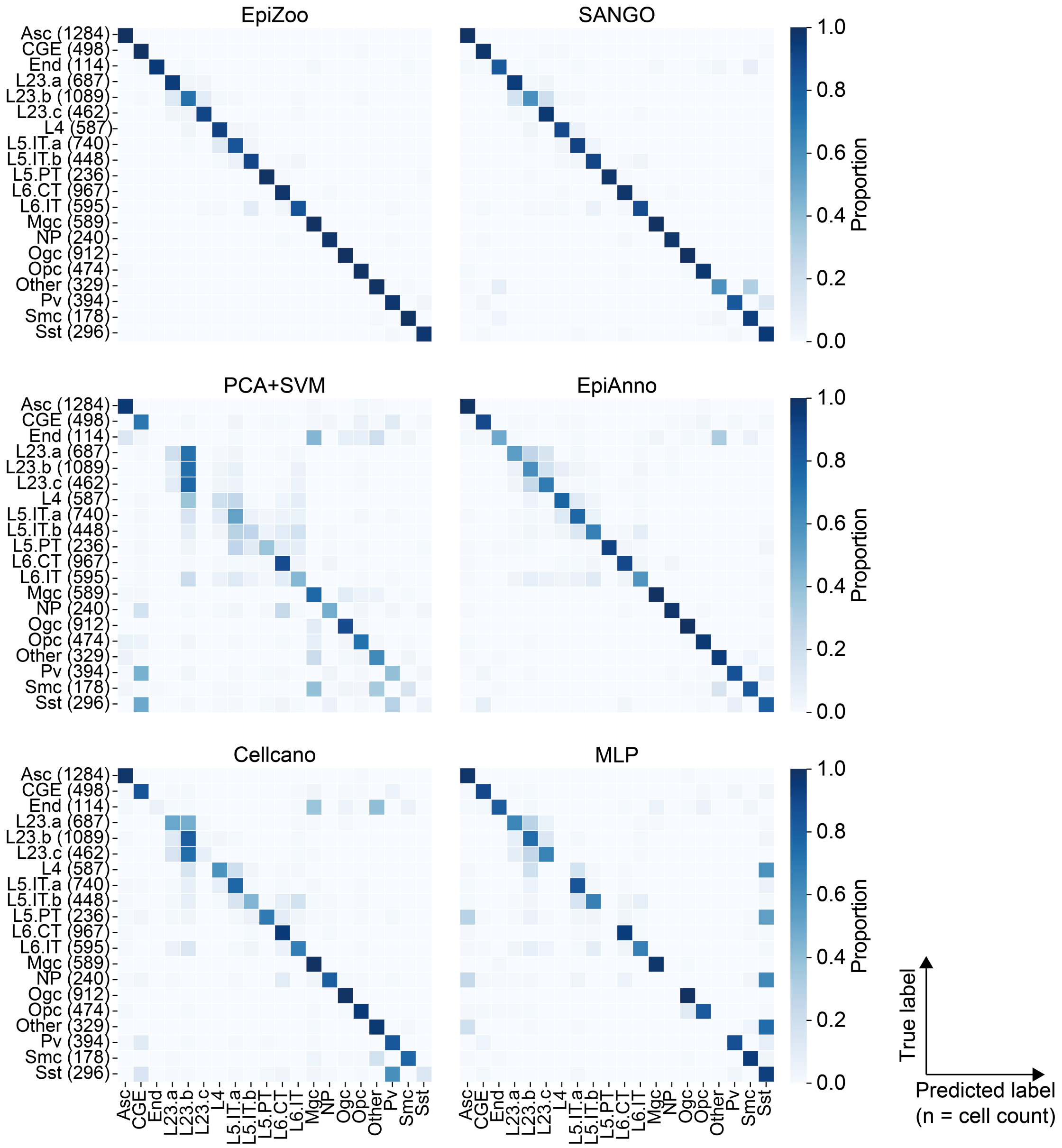


**Supplementary Figure 8.** Normalized confusion matrices of the Fang2021 dataset in cell-type annotation based on the intra-dataset five-fold cross-validation. Rows represent true cell types, and columns represent predicted cell types. Numbers in parentheses indicate the number of cells in each cell type.

### Supplementary Figure 9

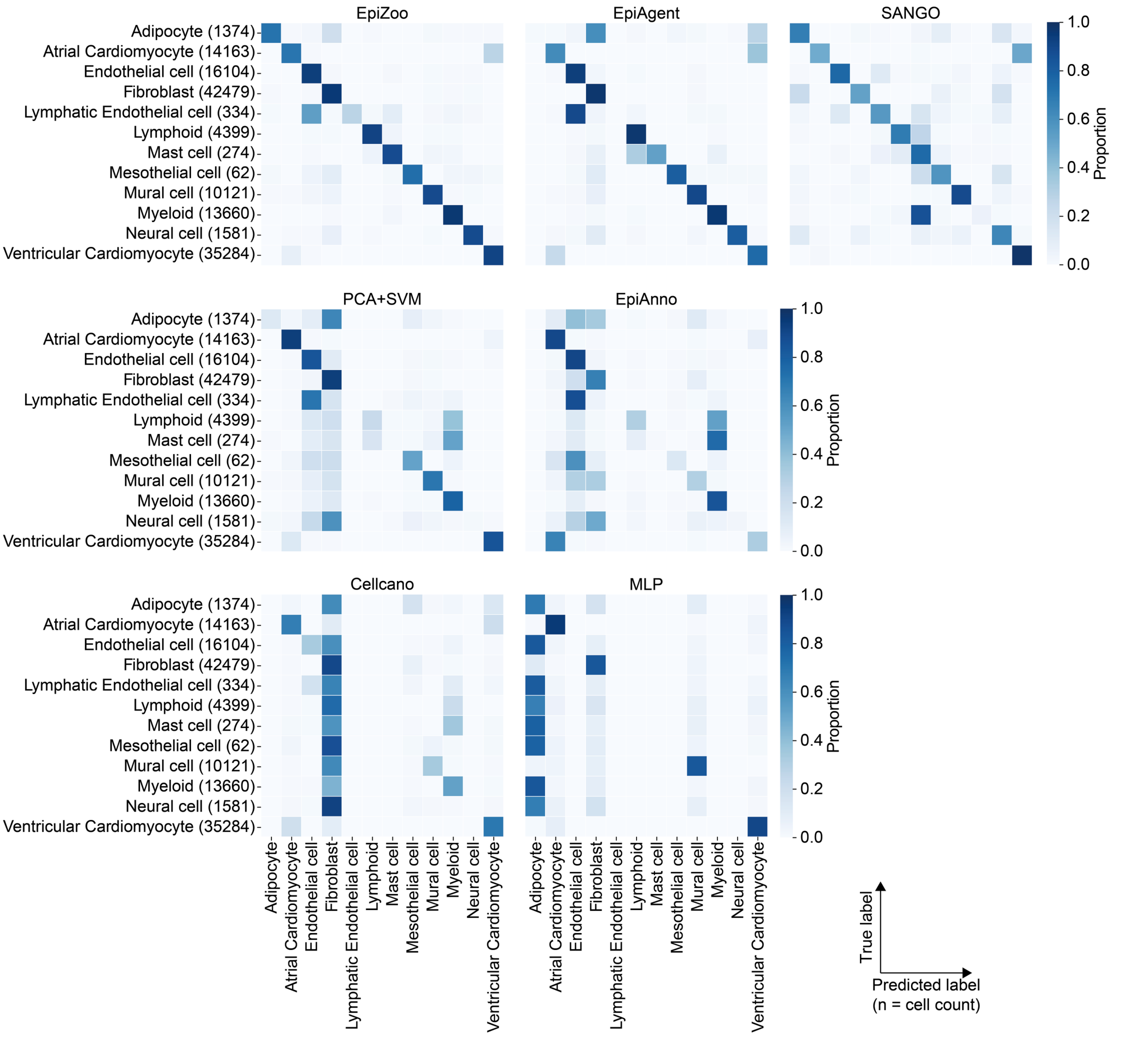


**Supplementary Figure 9.** Normalized confusion matrices of the Kanemaru2023 dataset in cell-type annotation based on the inter-dataset validation. Rows represent true cell types, and columns represent predicted cell types. Numbers in parentheses indicate the number of cells in each cell type.

**Supplementary Figure 10**

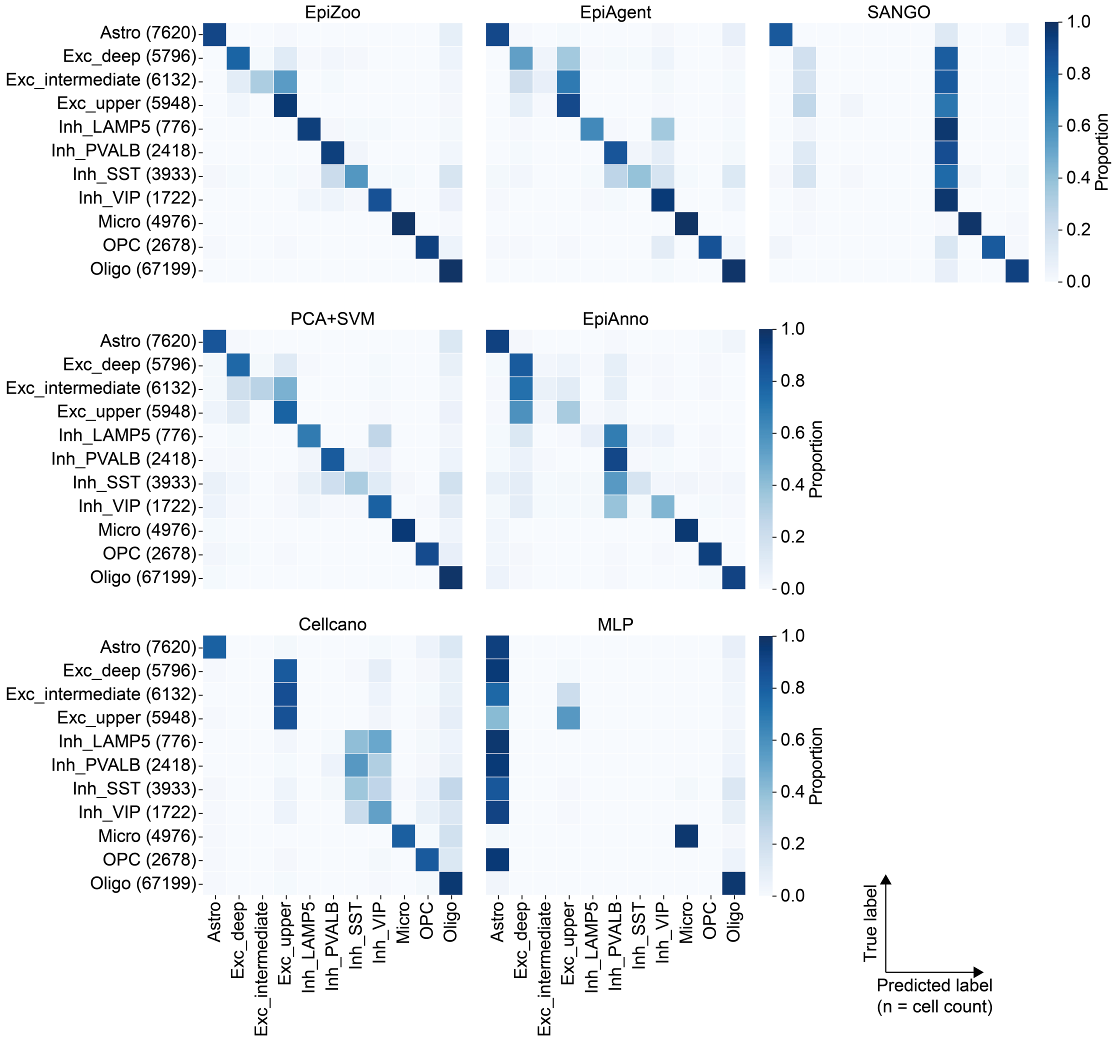


**Supplementary Figure 10.** Normalized confusion matrices of the Li2023b dataset in cell-type annotation based on the inter-dataset validation. Rows represent true cell types, and columns represent predicted cell types. Numbers in parentheses indicate the number of cells in each cell type.

### Supplementary Figure 11

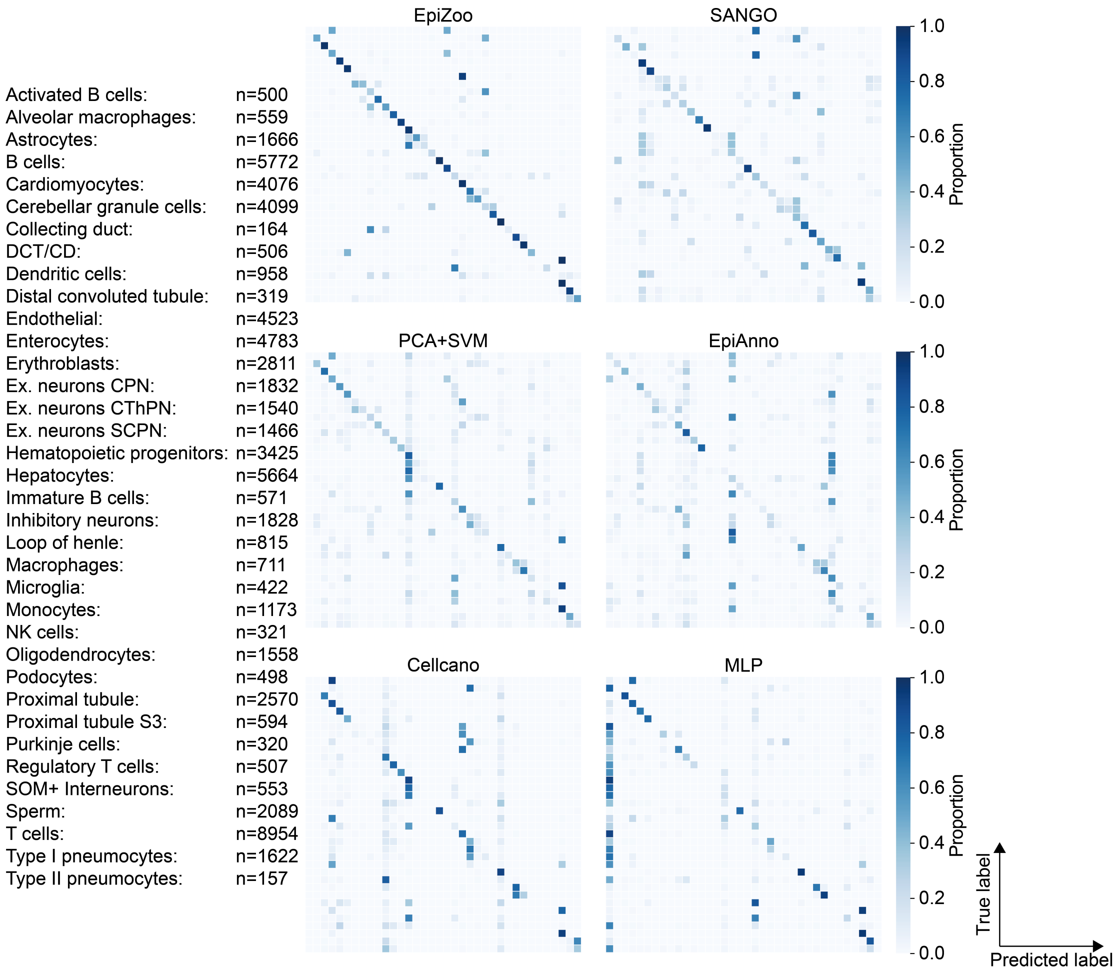


**Supplementary Figure 11.** Normalized confusion matrices of the Cusanovich2018 dataset in cell-type annotation based on the inter-dataset validation. Rows represent true cell types, and columns represent predicted cell types. Cell-type labels and corresponding cell counts shown on the left, aligning with the matrix rows.

### Supplementary Figure 12

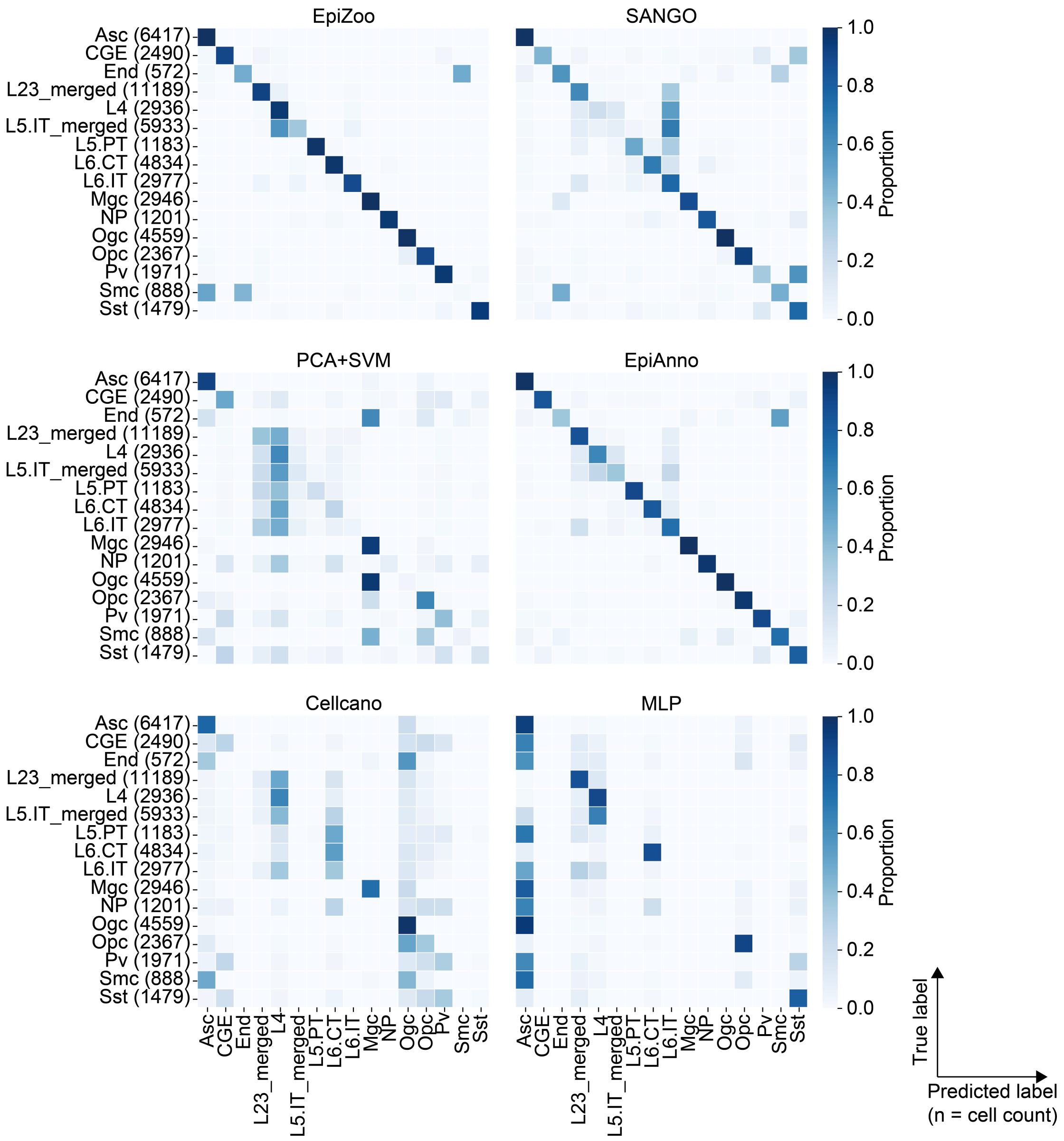


**Supplementary Figure 12.** Normalized confusion matrices of the Fang2021 dataset in cell-type annotation based on the inter-dataset validation. Rows represent true cell types, and columns represent predicted cell types. Numbers in parentheses indicate the number of cells in each cell type.

**Supplementary Figure 13**

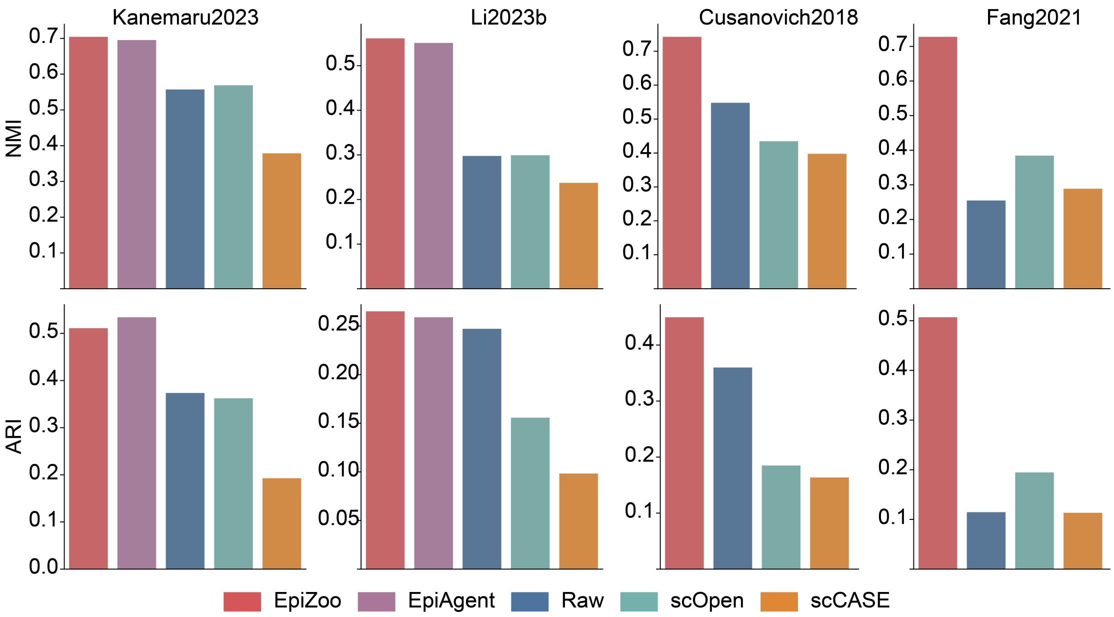


**Supplementary Figure 13.** Bar plots of NMI and ARI in data imputation across four datasets.

### Supplementary Figure 14

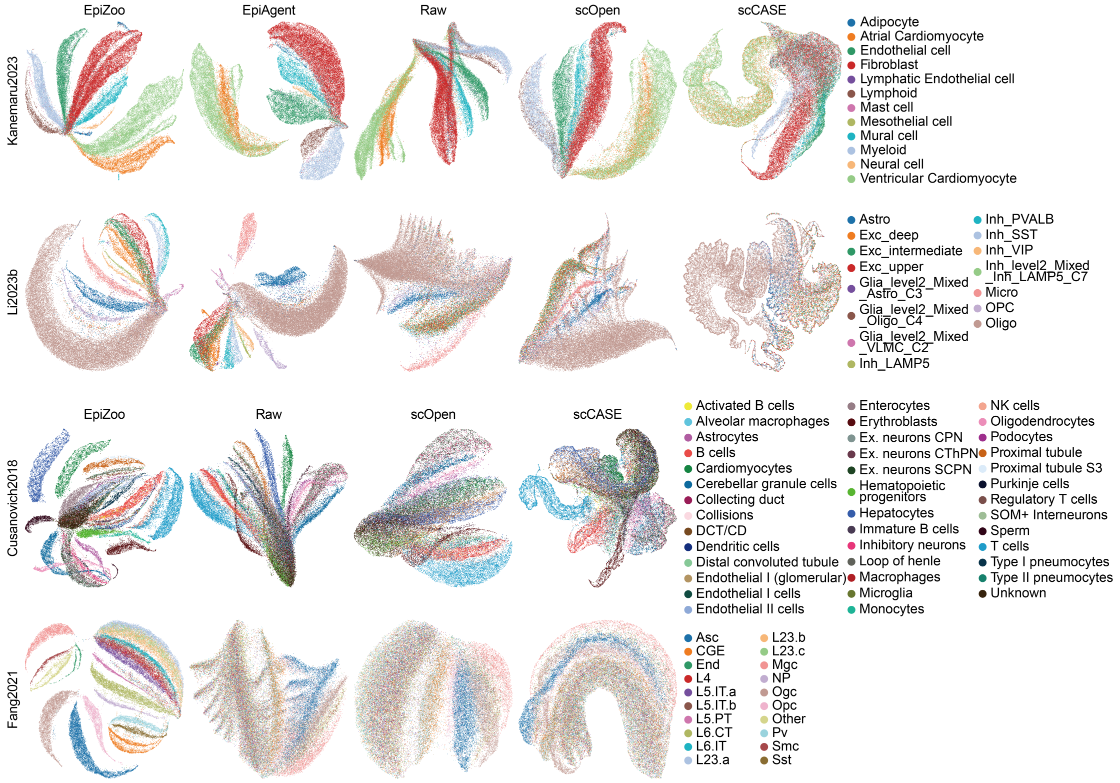


**Supplementary Figure 14.** UMAP visualizations of the imputed matrices derived from EpiZoo and baseline methods across four datasets.

### Supplementary Figure 15

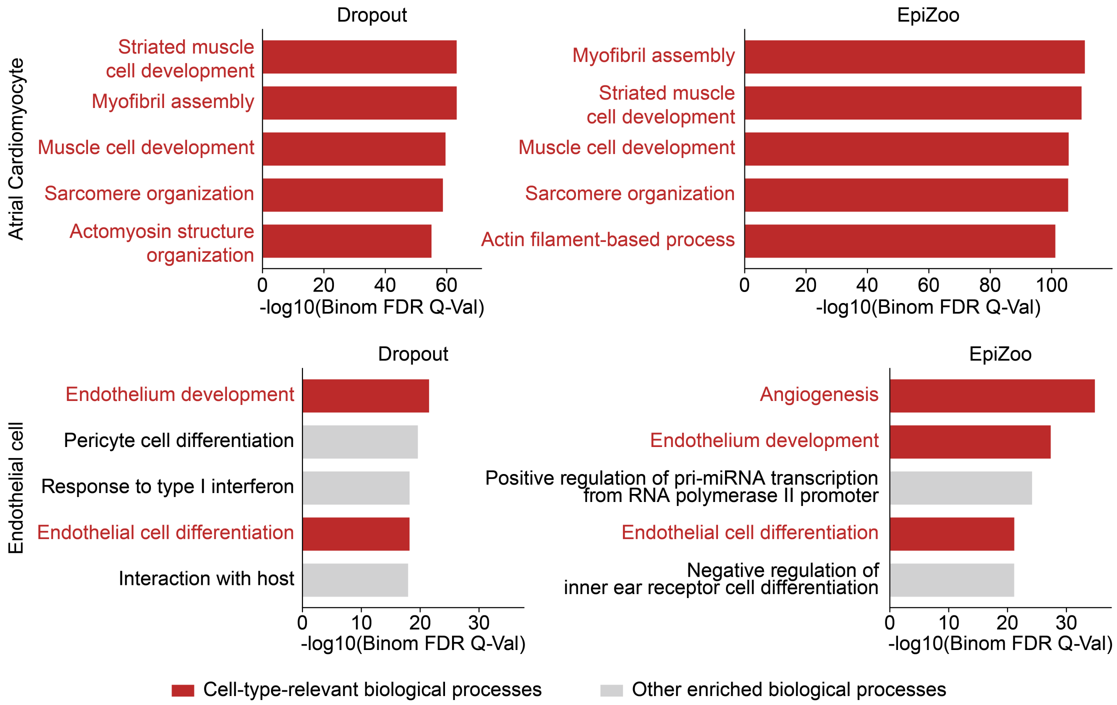


**Supplementary Figure 15.** Top five enriched biological processes identified by GREAT analysis for the top 5,000 cell-type-specific cCREs in atrial cardiomyocytes and endothelial cells from the Kanemaru2023 dataset.

**Supplementary Figure 16**

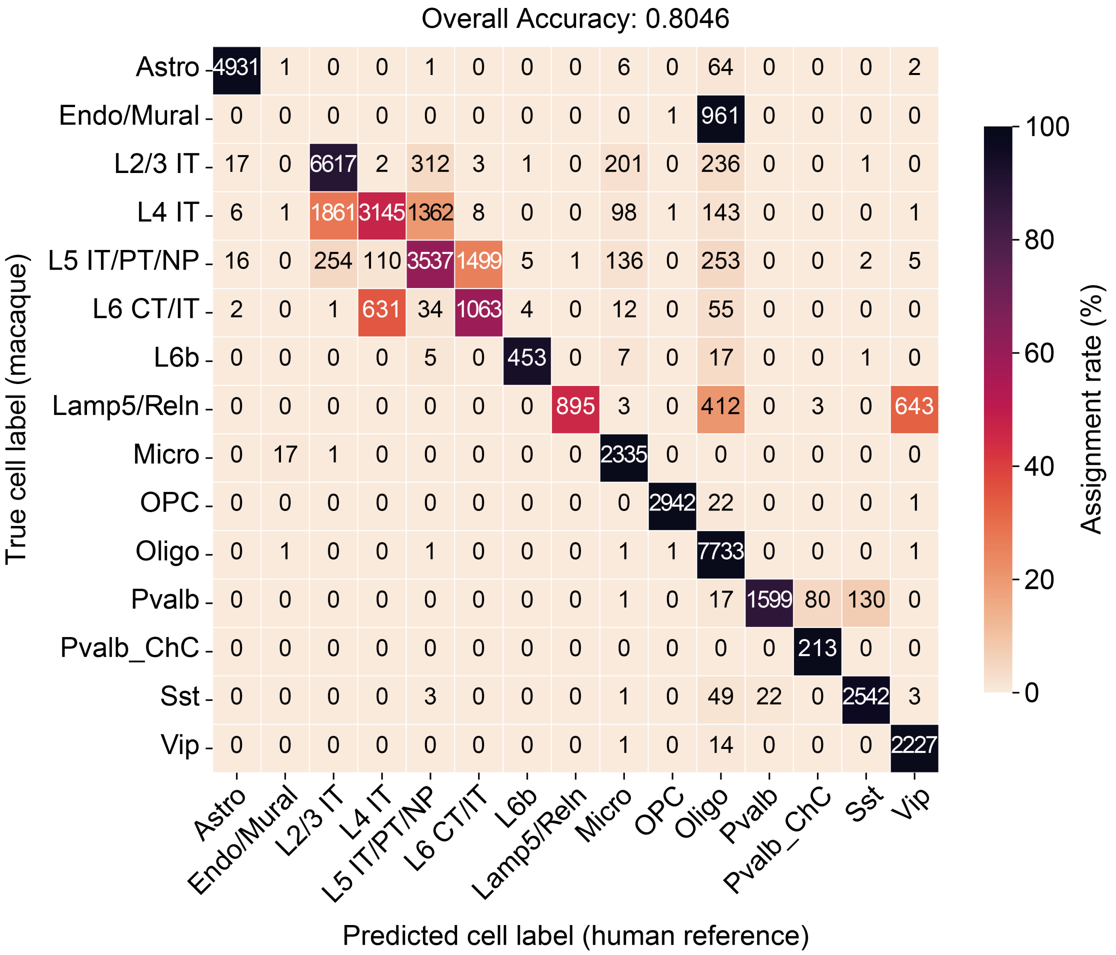


**Supplementary Figure 16.** Heatmap of zero-shot cross-species label transfer from human to macaque. Color scale indicates prediction accuracy; values represent cell counts.

### Supplementary Figure 17

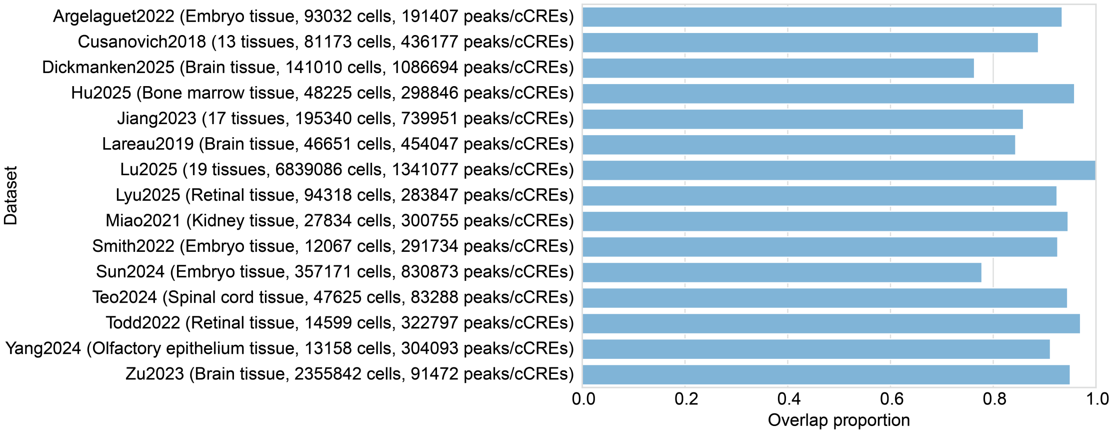


**Supplementary Figure 17.** The overlap proportion (%) between the peak/cCRE sets of each dataset and the cCRE set used by EpiZoo.
